# Insights on genomic profiles of drug resistance and virulence in a cohort of *Leishmania infantum* isolates from the Mediterranean area

**DOI:** 10.1101/2025.01.23.634476

**Authors:** Marina Carrasco-Martin, Joan Martí-Carreras, Marcel Gómez-Ponce, Maria Magdalena Alcover, Xavier Roura, Lluís Ferrer, Gad Baneth, Federica Bruno, Carmen Chicharro, Anabela Cordeiro-da-Silva, José Cristovão, Trentina Di Muccio, Carla Maia, Javier Moreno, Anabel Priego, Xavier Roca-Geronès, Nuno Santarem, Anna Vila Soriano, Fabrizio Vitale, Daniel Yasur-Landau, Olga Francino

**Affiliations:** Nano1Health S.L. (N1H), Edifici EUREKA, Parc de Recerca UAB, 08193 Bellaterra, Spain; Laboratori de Parasitologia, Departament de Biologia Sanitat i Mediambient, Facultat de Farmàcia i Ciències de l’Alimentació, Universitat de Barcelona, Av. Joan XXIII 27-31, 08028 Barcelona, Spain; Hospital Clínic Veterinari, Universitat Autònoma de Barcelona, 08193 Bellaterra, Spain; Departament de Medicina i Cirurgia Animals, Universitat Autònoma de Barcelona, 08193 Bellaterra, Spain; Laboratory for Zoonotic and Vector-Borne diseases - Koret School of Veterinary Medicine; Istituto Zooprofilattico Sperimentale della Sicilia; Instituto de Salud Carlos III; Instituto de Investigação e Inovação em Saúde (i3S), Universidade do Porto, 4200-135 Porto, Portugal and Departamento de Ciências Biológicas, Faculdade de Farmácia, Universidade do Porto, Rua de Jorge Viterbo Ferreira 228, 4050-313 Porto, Portugal; Global health and tropical medicine, LA-REAL, instituto de higiene e medicina tropical, universidade nova de Lisboa; Department of Infectious Diseases, Unit of Vector-Borne Diseases, Istituto Superiore di Sanità,Viale Regina Elena 299, 0161 Rome, Italy; Hospital Veterinario de la Universidad Católica de Valencia; SVGM, Molecular Genetics Veterinary Service, Universitat Autònoma de Barcelona, 08193 Bellaterra, Spain

**Keywords:** . *Leishmania infantum*, Copy Number Variation, Drug-resistance biomarkers, Virulence biomarkers, Canine Leishmaniosis, Human Leishmaniosis

## Abstract

**Background:** Drug-resistant strains of *Leishmania infantum* challenge the effectiveness of treatments for clinical leishmaniosis and may lead to more frequent relapses. Copy number variation (CNV) at specific genetic loci is associated with drug resistance and virulence, but information about its prevalence in endemic regions is limited. This study examines the drug resistance and virulence status of *Leishmania* strains in human and canine isolates from the Mediterranean region.

**Methods:** Forty-eight *Leishmania infantum* isolates were whole-genome sequenced with nanopore long reads, followed by *de novo* assembly. We analyzed chromosomal aneuploidies and gene copy number variation in loci linked to drug resistance and virulence in *Leishmania*, alongside the genomic structure and rearrangements responsible for these variations.

**Results:** Complete genomes were *de novo* assembled for 35 *L. infantum* isolates (22 from dogs and 13 from humans), revealing significant chromosomal variability. We assessed copy number variation for 22 potential biomarkers: 15 genes related to drug resistance to first-line drugs (METK for allopurinol; LdSMT for amphotericin B; AQP1 and H-locus for antimonials; LdMT, LdRos3, and MSL for miltefosine; PPM for paramomycin) and seven genes related to virulence (lipophosphoglycan and proteophosphoglycan biosynthesis, and the Lack protein). Drug-resistance biomarkers were identified in 80% of the isolates. Canine strains primarily showed resistance to allopurinol and antimonials, while human isolates exhibited a broader resistance spectrum, especially to antimonials and paromomycin. The co-occurrence of resistance biomarkers was common, especially for allopurinol and antimonial resistance. Distinct mechanisms underlie the observed copy number variations. Virulence-associated genes were less variable among isolates.

**Conclusions:** The prevalence of drug-resistance biomarkers in *Leishmania infantum* strains from the Mediterranean region, as revealed by this study, underscores the critical need for routine resistance surveillance in managing clinical leishmaniosis. These findings not only inform current clinical practice but also pave the way for more effective management strategies in the future.

## Background

Treatment failure in canine (CanL) and human zoonotic visceral leishmaniosis (HumL) might be influenced by the parasite (inherent virulence, drug resistance acquisition), and factors in the host (i.e., immunity) [1]. *Leishmania* (*L.) infantum* is the only autochthonous *Leishmania* species in the Mediterranean and peri-Mediterranean regions, causing both HumL and CanL. In these regions, WHO reported 8,367 cases of HumL over a 15-year period (2005-2020) [2] and CanL affects approximately 2.5 million dogs annually [3], both caused by *L. infantum*. The inherent virulence of each *Leishmania* strain depends on the parasite’s ability to adhere to and enter neutrophils and macrophages, protect against host factors, and prevent lysosome fusion. Thus, the parasite has evolved significant membrane-bound or secreted virulence factors, such as the glycocalyx molecules lipophosphoglycan (LPG) and proteophosphoglycan (PPG) [4].

For HumL in the Mediterranean region, liposomal amphotericin B (first registered in 1996 and exclusive to HumL use) is now the treatment of choice, replacing pentavalent antimonials (sodium stibogluconate and meglumine antimoniate) as the first-line drug, which had been in use since the 1920s [5]. Other alternatives to pentavalent antimonials now include miltefosine (first registered in India in 2002 [6] but with efficacy not proven in the treatment of European VL) and paromomycin (incorporated in 2006) [7]. The standard treatment for CanL combines leishmanicidal drugs, such as meglumine antimoniate or miltefosine (approved in Europe in 2007), with the off-label leishmaniostatic drug allopurinol, administered for long-term (6-12 months) [8].

The increasing emergence of antileishmanial drug-resistant *L. infantum* isolates is becoming a severe challenge to human and veterinary medicine, particularly in endemic regions [1, 9–11], rendering current antileishmanial drugs less effective. This situation requires increases in dose or prolonged treatment times, which can lead to associated side effects (e.g., nephrotoxicity). Resistant *Leishmania* strains to antimonials [12], allopurinol [13, 14], and miltefosine and amphotericin B [15] have been identified in infected dogs after treatment, as well as in humans [16]. These clinical cases underscore the role of drug resistance in disease relapse and reinforce the importance of antileishmanial-resistant *Leishmania* characterization to design effective disease control strategies aimed at informing therapeutic decision-making to reduce complications and treatment failure due to drug resistance.

Thus far, numerous studies have reported genomic information regarding gene copy number variation (CNV) associated with drug resistance and virulence. For instance, amplification into circular [17] or linear [18] minichromosomes of the multidrug-resistance protein A gene (*mrpa*) are associated with trivalent antimonial resistance, the active form resulting from the reduction of the pentavalent antimonial prodrug. A reduction in *L. donovani* miltefosine transporter gene (*LdMT*) decreases flippase activity diminishing miltefosine accumulation [19] and a reduction in *metk* CN reduces allopurinol sensitivity [20], respectively. Current evidence indicates that *Leishmania* strains deficient in LPG and PPG synthesis genes, as well as those lacking *lack* coding genes, result in attenuated virulence and non-viable parasites, respectively [4, 21–24].

In this context, this study aimed to describe and characterize the status of drug resistance and virulence biomarkers in a population of *L. infantum* isolated from the Mediterranean endemic area. Thus, we conducted a comprehensive genomic analysis of the *de novo* assembled genomes from *L. infantum* using Nanopore long reads whole-genome sequencing (WGS). Additionally, we defined a panel of 15 drug-resistance biomarkers for the most common treatments for leishmaniosis, along with 7 virulence biomarkers, and screened the cohort of 35 *L. infantum* isolates with LeishGenApp [25] to describe the state of drug resistance and provide relevant information for optimizing treatment selection for CanL and HumL.

## Methods

### Samples

Sixty *L. infantum* samples from promastigote cultures with parasite loads of 10^6^-10^8^ parasites/mL or *Leishmania* DNA were collected from reference collections or grown from clinical samples. Additionally, ATCC Control DNA was obtained from reference *L. infantum* strain Nicolle (ATCC 50134D, MHOM/TN/80/IPT-1). Out of 60 *L. infantum* samples, 48 were whole genome sequenced, and 35 isolates exceeded the quality threshold for CNV analysis (see below). These 35 isolates were obtained from different hosts and locations: 22 from dogs, 13 from humans, in Italy (Sicily 15), Spain (13), Portugal (4), Israel (2), and Tunis (1; the ATCC MHOM/TN/80/IPT-1 Control DNA).

The cultured samples were propagated in different media depending on their institution of origin: M199 medium, NNN medium cryopreserved in RMPI medium with 20% fetal bovine serum (FBS) and 10% DMSO, Evans’ modified Tobie’s medium, or Schneider medium (S0146-500ML, Sigma-Aldrich, St. Louis, MO, USA) supplemented with 20% FBS 1% human urine, and 25 mg/mL gentamicin, corresponding to cultures grown in institutions from Israel, Portugal, Sicily or Spain, respectively.

### DNA extraction and Nanopore Sequencing

DNA was extracted using the ZymoBIOMICS DNA Miniprep Kit (D4300; Zymo Research Corporation; Los Angeles, CA, USA). DNA quality and quantity were determined using a NanoDrop 2000 Spectrophotometer and a Qubit dsDNA HS Assay Kit (Fisher Scientific). Nanopore sequencing libraries with up to 2 barcoded samples per sequencing run were prepared using the Rapid Barcoding Sequencing Kit (SQK-RBK004; Oxford Nanopore Technologies, ONT) and loaded in a MinION FLO-MIN106 v9.4.1 flowcell or using the Native Barcoding Kit 24 V14 (SQK-NBD114.24; Oxford Nanopore Technologies, ONT) into a MinION FLO-MIN114 v.10.4.1 flowcell, respectively. Sequencing was performed on a GridION (GridION, ONT, Oxford, UK) for 48h. The fast5 files were basecalled and demultiplexed, and the adapters were trimmed using Guppy (v6.1.2-v7.1.4; “SUP” model) or Dorado (v7.2.13; “SUP” model).

### Chromosomal somy and gene copy number variation analysis

Chromosomal somy and gene copy number variation (CNV) were determined using the LeishGenApp v.2.2.0 analysis platform (Nano1Health S.L., Barcelona, Spain), as previously described (Martí-Carreras et al., 2022). Briefly, uncorrected nanopore reads were mapped against *L. infantum* reference (*L. infantum* JPCM5 v2/2018, http://leish-esp.cbm.uam.es/l_infantum_ downloads.html, accessed during 1 April 2022) [26] using LRA v1.3.7.2 [27]. The read alignments were processed by SAMTools v1.19.2 [28].

The detection of chromosomal somy and copy number variations (CNV) was done using CNVkit v0.9.10.1 [29]. Briefly, whole-genome sequencing (WGS) data were divided into consecutive bins of 10 kb and 1 kb for somy and CNV detection, respectively. For each isolate, a flat reference was generated, assigning a neutral state (log2 = 0.0) to all bins and incorporating GC content and RepeatMasker annotations from the reference genome to enable bias correction (- gc - rmask -o reference.cnn). The observed read depths were calculated with coverage function and normalized with the fix command, which median-centers, bias-correct the data, and subtracts the expected log2 values from the reference.cnn file, yielding to .cnr files containing corrected bin-level log2 ratios with associated weights. The bin-level ratios were then segmented, producing .cns files that summarize contiguous genomic regions with homogeneous log2 ratios. Finally, the robustness of each segment was evaluated with segmetrics, which reports residual variability and confidence intervals for segment means, providing additional support for CNV calls. Therefore, a genomic coordinate of an isolate is considered as one with "CNV" if it shows a significant change in median-centered, bias-corrected log2 ratio in a window length of 1 kb compared to its flat reference at the same given genomic coordinate (Additional file 2: Table S2).

The chromosome copy number was determined by calculating the median of all the log2 ratio values derived for all 10-kb bin measurementst of each chromosome. The somy values were defined as S < 1.5, 1.5 < S < 2.5, 2.5 < S < 3.5, and 3.5 < S < 4.5 for for monosomy, disomy, trisomy and tetrasomy, as described in Patino et al., 2019.

Gene copy number was calculated by the logarithmic transformation of the log_2_ ratio measurements corresponding to genomic coordinates of the 15 drug target and transporter genes and 7 virulence genes. A threshold for biomarker detection of ±0.50 from the sensitive-linked copy number (2, 3, or 4 copies) was used (see Table 1), considering a 5% natural polymorphism within the population [31], and the LeishGenApp-associated variability (±0.38, data not shown). Samples with less than 10X coverage for chromosomes with resistance biomarker genes were excluded from the analysis.

**Table 1.**
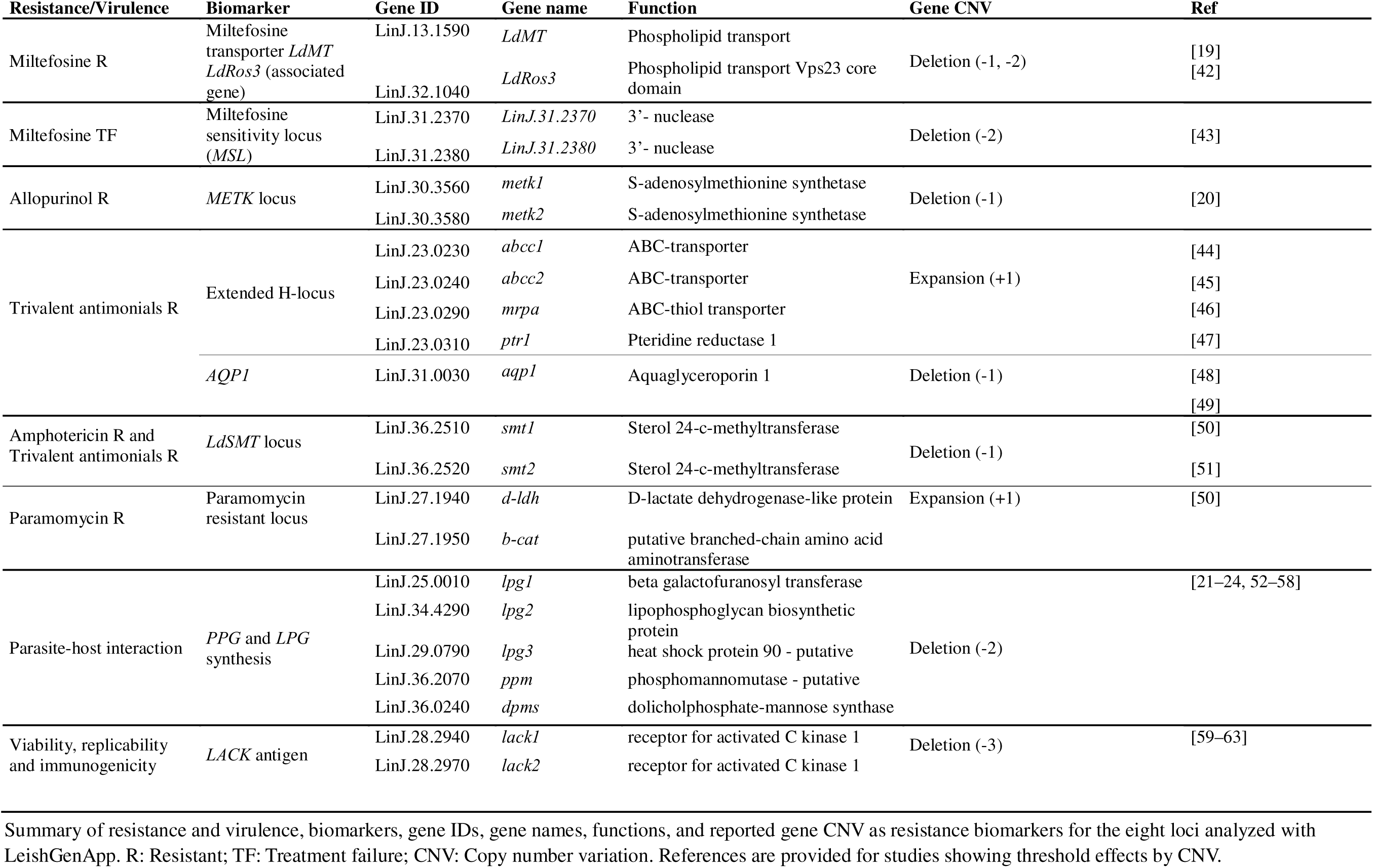
CNVs linked to drug resistance and virulence in *L. infantum* treatment.

*METK, LdMT,* and *mrpa* gene copy numbers were verified by using LeishGenR^TM^ (NANO1HEALTH SL (Barcelona, Spain)) based on a molecular quantitative genomic approach.

### Genome *de novo* assembly and annotation

Genomes were *de novo* assembled using N1H-Assemblatron v2.1.1 (Nano1Health SL, Barcelona, Spain) discarding reads with Phred-score lower than 10 by Filtlong v0.2.1. Data summary statistics were obtained using Nanoplot v.1.42.0 [32]. Flye v2.9.1-v2.9.3 [33] was used for *de novo* assembly (-nano-hq) and contigs were polished using Medaka v1.7.2-2.0.0 (https://github.com/nanoporetech/medaka) (-m r941_min_sup_g507). Genome completeness was assessed using Busco v5.4.45-v5.8.0 [34]. Genomes were annotated using online Companion platform v2.2.8 [35].

### Identification of repeated sequences in the drug resistance associated loci of *L. infantum*

Unknown and described repeats were screened within the extended H-locus regions: LinJ.23:59772-106565, the *METK* locus: LinJ.30:1280005-1294327, and the SMT locus: LinJ.36:952046-960534 in the *L. infantum* reference genome JPCM5 (GenBank accession number GCA_000002875.2). Locations of the repeated sequences were identified with nucmer (-l 50) [36] and blastn (part of blast+) [37]. We filtered repeated sequences longer than 0.5 kb. The repeated sequences observed in the reference genome JPCM5 were identified in the *de novo* assemblies of our 35 isolates using the same alignment tools. IGV v2.16.1 [38] was used to visualize, analyze, and describe the structure of regions with repeated sequences.

### Data analysis and visualization

A linear regression model assessed whether mid-term culturing significantly explains aneuploidy levels. The model included one factor with two levels: clinical isolates (<10 passages, n=10) and culture lines (10–33 passages and/or subjected to a freezing step, n=9). See Table 2 for in detail sample information. Assumptions of homoscedasticity (Bartlett’s test: K² = 0.345, p = 0.556) and residual normality (Shapiro-Wilk: W = 0.967, p = 0.229) were met. Wilson’s method was used to calculate 95% confidence intervals (95% CIs) for drug resistance biomarker prevalence. Data analysis and visualization were performed using R-packages, including tidyverse [39], and BioCircos [40], using R studio interface running R version 4.4.1.

**Table 2.** Sample ID and metadata used in this study. Thirty-five L. infantum samples were used in this study. Columns represent the relation with their original host, collection year, geographical location and detected resistance biomarkers by using LeishGenApp.

| Sample ID | Host species | Collection Year | Location | Passages | Resistance biomarkers |  |  |  |  |
| --- | --- | --- | --- | --- | --- | --- | --- | --- | --- |
|  |  |  |  |  | Allopurinol | Amphotericin B | Antimonials | Miltefosine | Paromomycin |
| MCAN/PT/2012/IMT397 | *Canis lupus familiaris* | 2012 | Portugal (Setúbal) | - | METK | - | - | - | - |
| MHOM/MA/67/ITMAP-263 | *Homo sapiens sapiens* | 2023 | Portugal | - | METK | - | *aqp1* | - | - |
| MHOM/MA/67/ITMAP-264 | *Homo sapiens sapiens* | 2023 | Portugal | - | - | *smt1* | *abcc1, abcc2, mrpa* | - | - |
| MHOM/MA/67/ITMAP-265 | *Homo sapiens sapiens* | 2023 | Portugal | - | - | - | *abcc1, abcc2, mrpa, aqp1* | - | - |
| LCAN/ES/1992/CATB011 | *Canis lupus familiaris* | 1992 | Spain (Catalonia) | - | - | - | *abcc1, abcc2, mrpa, ptr1, aqp1* | - | - |
| MCRI/ES/2006/CATB033 | *Canis lupus familiaris* | 2006 | Spain (Catalonia) | - | - | - | - | *LdMT* | - |
| MHOM/ES/2016/CATB101 | *Homo sapiens sapiens* | 2016 | Spain (Mallorca) | 11 | - | - | *aqp1* | - | *b-cat* |
| LCAN/ES/2020/CATB102 | *Canis lupus familiaris* | 2020 | Spain (Zaragoza) | 8 | - | - | - | - | *b-cat, d-ldh* |
| LCAN/ES/2020/CATB103 | *Canis lupus familiaris* | 2020 | Spain (Zaragoza) | 7 | - | - | *aqp1* | - | - |
| LCAN/ES/2022/CATB104 | *Canis lupus familiaris* | 2022 | Spain (Valencia) | 4 | - | - | - | - | - |
| LCAN/ES/2023/CATB105 | *Canis lupus familiaris* | 2023 | Spain (Barcelona) | 5 | - | - | - | - | - |
| LCAN/ES/2023/CATB108 | *Canis lupus familiaris* | 2023 | Spain (Barcelona) | 11 | - | - | - | - | - |
| LCAN/ES/2023/CATB109 | *Canis lupus familiaris* | 2023 | Spain (Barcelona) | 15 | - | - | - | - | - |
| LCAN/ES/2024/CATB110 | *Canis lupus familiaris* | 2024 | Spain (Valencia) | 5 | - | - | *aqp1* | - | - |
| MCAN/IL/2011/NT2 | *Canis lupus familiaris* | 2011 | Israel | - | METK | *smt1* | *abcc1, mrpa, ptr1, aqp1* | - | *b-cat* |
| MHOM/TN/80/IPT-1 | *Homo sapiens sapiens* | 1980 | Tunisia (Monastir) | - | METK | *smt1, smt2* | *abcc1, abcc2, mrpa, ptr1* | - | *d-ldh* |
| MHOM/ES/2022/LLM-2503 | *Homo sapiens sapiens* | 2022 | Spain (Zaragoza) | - | - | - | - | - | - |
| MCAN/IT/2011/8283 | *Canis lupus familiaris* | 2011 | Italy (Sicily, Palermo) | 11 | METK | - | - | - | - |
| MCAN/IT/2022/21SCIACCA | *Canis lupus familiaris* | 2022 | Italy (Sicily, Agrigento) | 6 | METK | - | *abcc1, abcc2, mrpa, ptr1, aqp1* | - | - |
| MHOM/IT/2017/1533U | *Homo sapiens sapiens* | 2017 | Italy (Sicily, Agrigento) | 6 | - | - | *aqp1* | - | *b-cat, d-ldh* |
| MCAN/IT/2018/4586 | *Canis lupus familiaris* | 2018 | Italy (Sicily, Palermo) | 20 | - | - | *aqp1* | - | - |
| MCAN/IT/2018/1380 | *Canis lupus familiaris* | 2018 | Italy (Sicily, Palermo) | 33 | METK | - | - | - | - |
| MHOM/IT/2008/31U | *Homo sapiens sapiens* | 2008 | Italy (Sicily, Palermo) | 7 | - | - | - | - | - |
| MCAN/IT/2020/1265 | *Canis lupus familiaris* | 2020 | Italy (Sicily, Palermo) | - | METK | - | *abcc2, mrpa* | - | - |
| MCAN/IT/2022/2885 | *Canis lupus familiaris* | 2022 | Italy (Sicily, Palermo) | 26 | METK | - | *aqp1* | - | *b-cat, d-ldh* |
| MCAN/IT/2020/4299 | *Canis lupus familiaris* | 2020 | Italy (Sicily, Palermo) | 10 | METK | - | - | - | - |
| MCAN/IT/2021/18017 | *Canis lupus familiaris* | 2021 | Italy (Sicily, Palermo) | 8 | METK | - | *abcc2* | - | *b-cat* |
| MCAN/IT/2019/3299 | *Canis lupus familiaris* | 2019 | Italy (Sicily, Palermo) | 19 | - | - | - | *LdMT* | - |
| MCAN/IT/2019/4549 | *Canis lupus familiaris* | 2019 | Italy (Sicily, Palermo) | 9 | METK | - | *abcc1, abcc2, mrpa, ptr1* | - | - |
| MHOM/ES/2022/LLM-2505 | *Homo sapiens sapiens* | 2022 | Spain (Toledo) | - | - | - | *aqp1* | - | - |
| MHOM/IT/90/ISS509 | *Homo sapiens sapiens* | 1990 | Italy (Sicily, Catania) | - | - | - | - | - | *b-cat, d-ldh* |
| MHOM/IT/91/ISS666 | *Homo sapiens sapiens* | 1991 | Italy (Sicily, Catania) | - | - | - | - | - | - |
| MHOM/IT/93/ISS828 | *Homo sapiens sapiens* | 1993 | Italy (Sicily, Catania) | - | - | - | - | *LdRos3* | *b-cat* |
| MHOM/ES/2021/LLL-2500 | *Homo sapiens sapiens* | 2021 | Spain (Madrid) | - | - | - | *abcc1, aqp1* | - | - |
| MCAN/IL/2011/NT6 | *Canis lupus familiaris* | 2011 | Israel | - | METK | - | *abcc1, ptr1* | - | - |

## Results

### Detection of aneuploidies and genome plasticity in *L. infantum* genome by nanopore-only whole-genome sequencing

To characterize the status of drug resistance and virulence biomarkers in *L. infantum* isolates, we collected 60 isolates from either dogs or humans within the Mediterranean basin, an area endemic to CanL and HumL. The isolates came from Italy (20), Portugal (20), Spain (13), and Israel (7), with titers ranging from 10^6^ to 10^8^ parasites/ml. Forty-three isolates with titers higher than 10^7^ parasites/ml were whole genome sequenced, with a N50 ranging between 6 and 9 kb. After quality filtering, 35 isolates—22 canine and 13 humans—with sufficient parasite load (>10D parasites/ml) and sequencing depth (average sequencing depth of 58X; genome coverage 97%) were retained for downstream analysis. Eight isolates were excluded due to insufficient depth across chromosomes (<10X). Data is available in the DDBJ/ENA/GenBank repository under the BioProject PRJNA881045 with the following accession numbers: JAZFNU000000000-JAZFNZ000000000; JAZFOA000000000-JAZFOZ000000000, and JAZFPA000000000-JAZFPC000000000 and, assembly metrics are in Additional file 1: Table S1.

The detailed analysis of the assembled *L. infantum* genomes reveals significant structural variability at the chromosomal levels, with just one canine isolate (LCAN/ES/2024/CATB110) exhibiting the expected karyotype—disomy across all chromosomes except for tetrasomy of chromosome 31 (Figure 1a), and high karyotypic diversity between isolates. Read-depth analysis reveals isolate-specific variations in some of the 36 chromosomes, showing either increases or decreases in coverage across the entire chromosome. For instance, chromosome 31 consistently shows a two-fold increase (Figure 1a). Based on chromosomal somy, three groups were identified: (i) stable disomy in at least 95% of isolates for chromosomes 7, 14, 18, 19, 27, 28, 34, and 36; (ii) consistent tetrasomy of chromosome 31 in 88% of isolates (SD = 0.29) as expected for *L. infantum* [41]; and (iii) variable somy observed in the remaining 27 chromosomes. Our approach revealed that trisomy was common for chromosomes 8, 20, and 23 (Figure 1b). However, high inter-isolate variability was noted for chromosomes 8 and 23, with a standard deviation (SD) ≥ 0.87 and a notable difference between the mean (2.80) and median somy values (2.48) of chr. 23. Notably, aneuploidies in chromosomes 23 and 31 are of particular interest, as they are coding seven of the 15 genes whose CNV are linked to drug resistance (Table 1).

**Figure 1.**
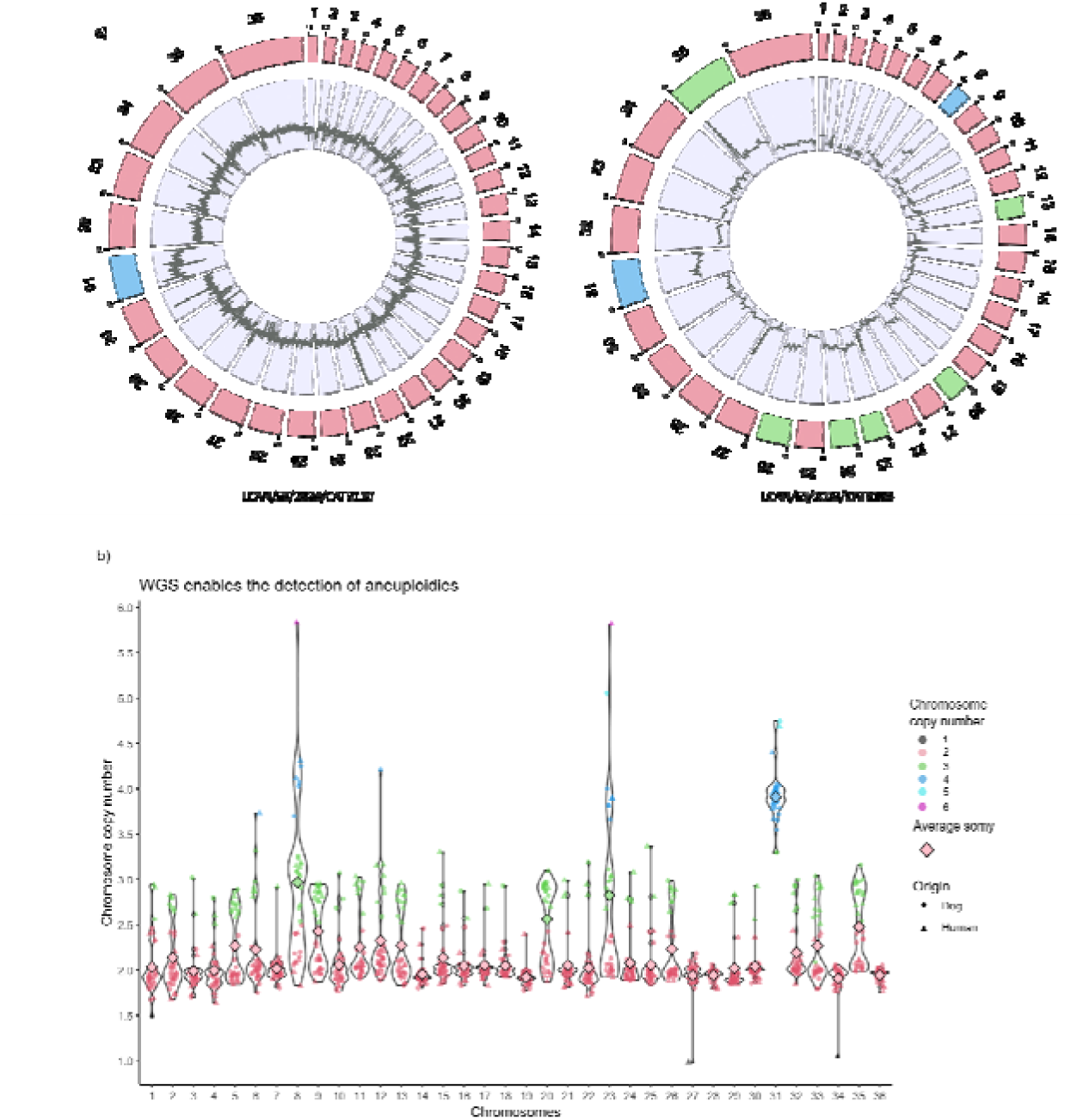
Aneuploidies and mosaicism in the genomes of *Leishmania infantum* isolates. a) Circular representation of whole-genome coverage by BioCircos. The outer circle represents the 36 chromosomes, and the inner circle represents the coverage along each chromosome. Sample LCAN/ES/2024/CATB110 has the expected disomic background except for tetrasomic chromosome 31. Sample LCAN/ES/2023/CATB108 shows aneuploidy for chromosomes 13, 20, 23, 24, 26 and 35 (trisomy) and 8 and 31 (tetrasomy). b) Violin plots showing the chromosome copy number for each of the 35 *L. infantum* chromosomes across all samples. The median somy for each chromosome, calculated as the median copy number across the 35-sample cohort, is shown by the bigger rhombus.

Overall, polyploidy was common, with 12 isolates (33%) having ≥10 chromosomes with somy >2, while monosomy was rare and restricted to three events across two isolates. Intermediate somy values, non-integer copy numbers, were frequently observed, consistent with mosaic aneuploidy (Figure 1b).

A comparison of clinical isolates (≤10 in vitro passages, n = 10) with mid-passage cultures (10–33 passages, n = 9) did not yield significant differences in aneuploidy levels resulting from mid-term culturing (F = 2.383, R² = 0.122, p = 0.141). Nevertheless, higher aneuploidy was observed in the long-passaged reference strain *L. infantum* IPT-1 (10 aneuploid chromosomes) and sample MCAN/IT/2020/1265 (68 passages; 11 aneuploid chromosomes), which were excluded from the analysis as they are considered clonal lines because of their extensive passage history (Table 2).

### Selection of CNV as drug resistance and virulence biomarkers in *L. infantum*

We selected CNV in genes present at low copy number (1–2 copies) in the genome that have previously been linked to drug resistance and virulence in *Leishmania infantum* (see Table 1). We identified 22 CNV as potential biomarkers in *L. infantum*: 15 related to drug resistance and seven related to virulence. The drug resistance panel includes genes from eight loci: METK, LdSMT, AQP1, LdMT, LdRos3, MSL, PPM, and the extended H-locus (38 kb), which contains *abcc1*, *abcc2*, *mrpa*, and *ptr1*. CNV in these genes are associated with responses to the five main drugs used for treating leishmaniasis: allopurinol, amphotericin B, pentavalent antimonial, miltefosine, and paramomycin. The seven virulence-linked CNVs are involved in early infection steps (i.e. neutrophils phagocytosis, entry into macrophages, and resistance to complement-mediated lysis) (5/7) and in aspects of parasite viability, replication, and immunogenicity (2/7) (see Table 1). This panel of 22 CNV biomarkers forms the core of LeishGenApp (25) and was screened against the 35 *L. infantum* isolates.

### *L. infantum* isolates from dogs and humans show distinct drug-resistance biomarker status while mostly preserving virulence

Across the *L. infantum* mediterranean cohort, 80% (28/35) of isolates harbored at least one drug-resistance biomarker, with an average of 2.4 biomarkers per isolate (SD = 2.2), ranging from 8 (MHOM/TN/80/IPT-1) to 0 (7 isolates) (Figures 2a and 2c; Table 2; Additional file 2: Table S2).

**Figure 2.**
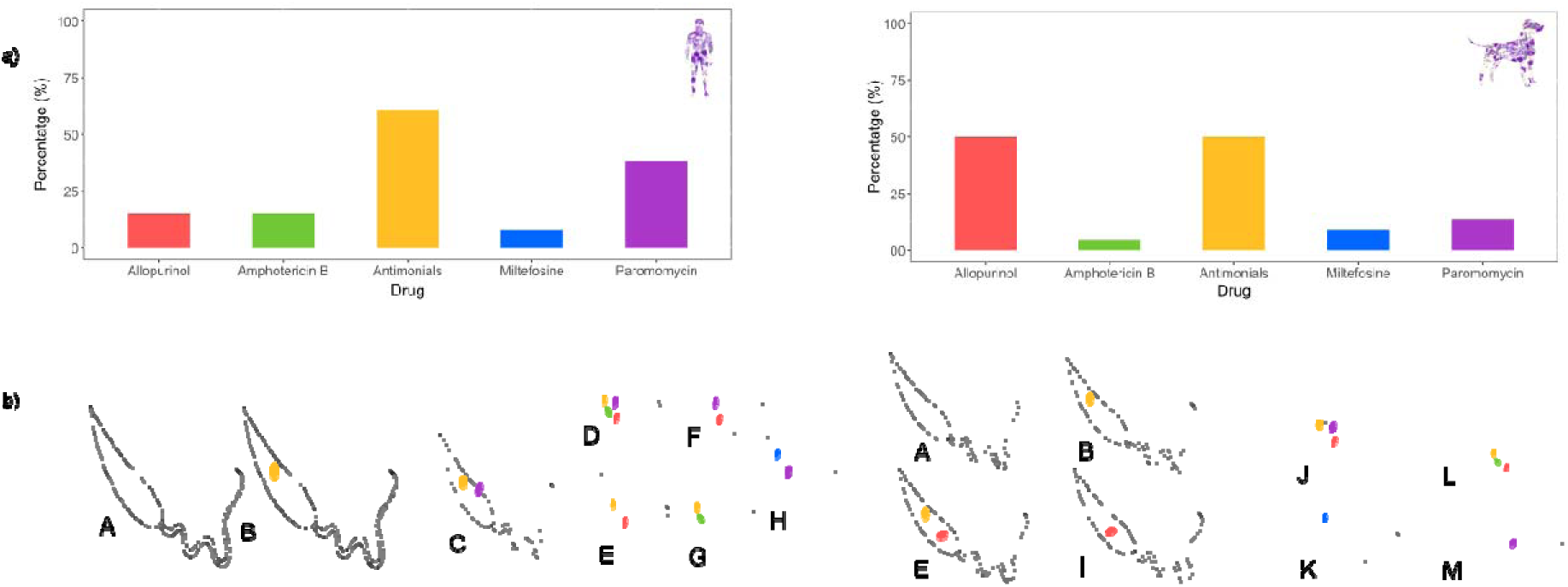
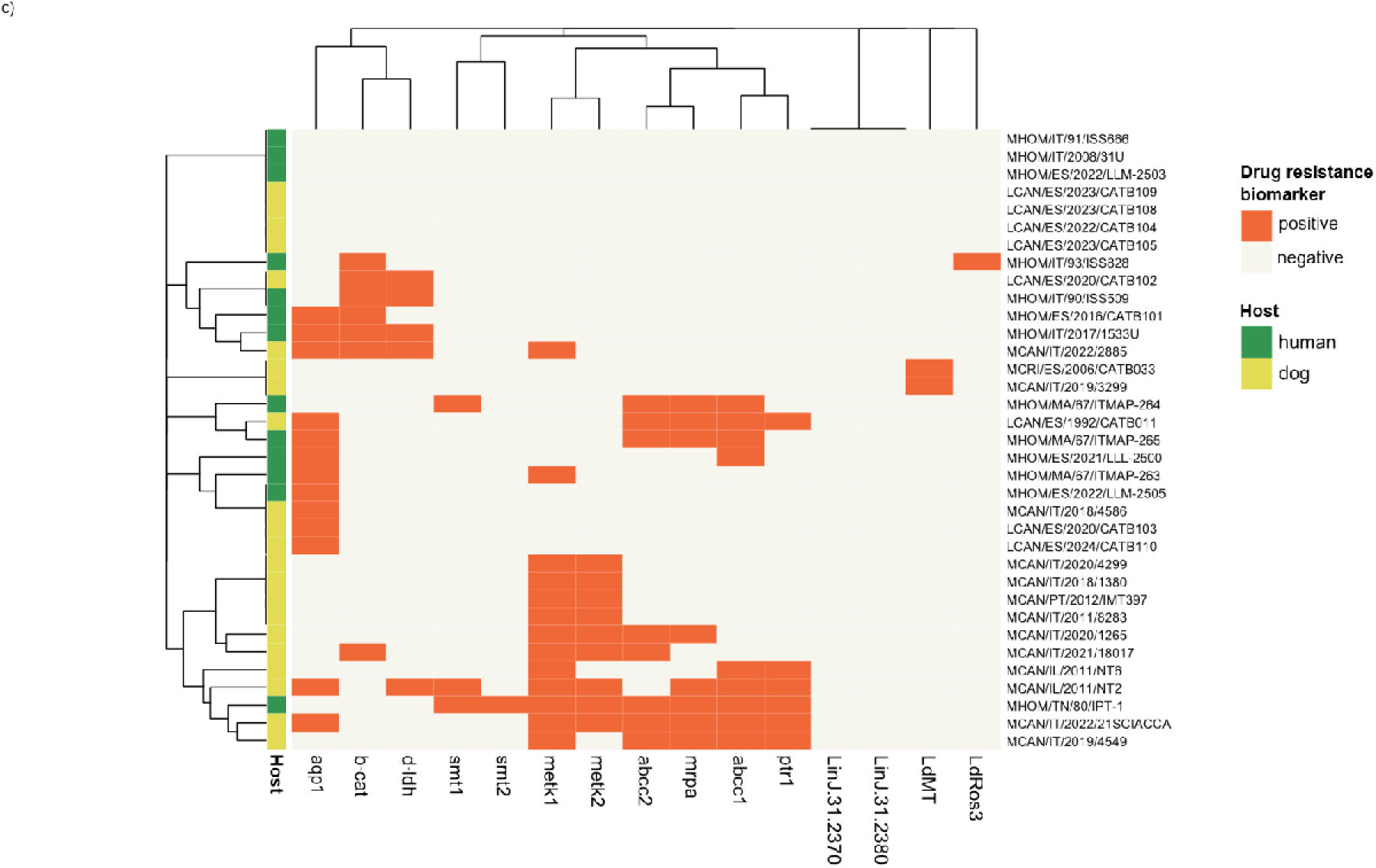

The prevalence of drug-resistance biomarkers to the five first-line drugs varied across the cohort. Antimonial resistance was the most common (54.2%), followed by allopurinol resistance (37.1%), paromomycin resistance (22.8%), and resistance biomarkers for both miltefosine and amphotericin B (8.5% each) (See Table 3). Notably, antimonial resistance biomarkers were frequently found in CanL and HumL isolates (61% in humans; 50% in dogs). Although host-specific patterns seem to emerge, with 50% of canine isolates testing positive for allopurinol resistance biomarkers, compared to 15% of human isolates, no statistical difference was found between the host groups (P = 0.07) (See Table 3). Meanwhile, resistance to paromomycin (38% vs. 14%) and amphotericin B (15% vs. 4.5%) was more common in human isolates. Hierarchical clustering suggested a tendency for resistance profiles to cluster by host (F = 1.87, P = 0.088), with a distinct canine cluster characterized by the presence of the *METK* deletion (see Results section below), a known marker for allopurinol resistance, and, in some cases, the presence of H-locus amplification, a marker for antimonial resistance, also found in the high-passage reference strain IPT-1 (Figure 2c).

**Table 3.** Prevalence of drug-resistance biomarkers in *L. infantum* and comparison of each antileishmanial resistance prevalence between host groups.

| Variable/ Category | Positive for drug-resistance biomarker(s)<br>n (%; 95% IC) | P-value |
| --- | --- | --- |
| **5 antileishmanial drugs** | 28 (**80**, 64.1 – 89.9) | $P = 1$ |
| Human | 18 (**81**, 61.4 – 92.6) |  |
| Dog | 10 (**76**, 49.7 – 91.8) |  |
| **Allopurinol** | 13 (**37.1**, 23.1 – 53.6) | $P = 0.07$ |
| Human | 2 (**15.4**, 4.3 – 42.2) |  |
| Dog | 11 (**50**, 30.7 – 69.3) |  |
| **Amphotericin B** | 3 (**8.5**, 2.9 – 22.3) | $P = 0.54$ |
| Human | 2 (**15.4**, 4.3 – 42.2) |  |
| Dog | 1 (**4.5**, 0.8 – 21.8) |  |
| **Antimonials** | 19 (**54.2**, 38.1 – 69.5) | $P = 0.72$ |
| Human | 8 (**61.5**, 35.5 – 82.2) |  |
| Dog | 11 (**50**, 30.7 – 69.3) |  |
| **Miltefosine** | 3 (**8.5**, 2.9 – 22.3) | $P = 1$ |
| Human | 1 (**7.7**, 1.4 – 33.3) |  |
| Dog | 2 (**9.1**, 2.5 – 27.8) |  |
| **Paromomycin** | 8 (**22.8**, 12 – 39) | $P = 0.43$ |
| Human | 5 (**38.4**, 17.7 – 64.4) |  |
| Dog | 3 (**13.6**, 4.7 – 33.3) |  |

We assessed the drug-resistance biomarker status for each isolate and categorized them into profiles to analyze the similarities and differences between CanL (22) and HumL (13). Thirteen distinct resistance profiles (labeled A–M) were identified (Figure 2b). Three profiles (A: no resistance; B: antimonial resistance; and E: dual resistance to antimonials and allopurinol) were shared between dogs and humans, and were the most common, accounting for 54% of all isolates. The remaining ten profiles were host-specific, with five exclusive to dogs and five to humans (Figure 2b), which aligns well with the clustering results.

The eight HumL profiles were distributed heterogeneously (mean = 1.62, sd = 0.85, CV = 53%). The most prevalent profiles were negative (A; 23%) and those exhibiting resistance to antimonials (B; 23%) (Figure 2b). Likewise, CanL profiles exhibited a heterogeneous distribution (mean = 2.75, sd = 1.29, CV = 47%). However, they appeared to be more clustered, with 73% of the isolates distributed across four profiles: A (negative for drug-resistance biomarkers), B (antimonials), E (antimonials and allopurinol), and I (allopurinol), each encompassing four isolates. It is noteworthy that 50% of CanL isolates were associated with resistance to allopurinol (Figure 2b). Moreoever, biomarkers for resistance to allopurinol and antimonials co-occurred in 32% of isolates. These biomarkers consistently clustered together in the dendrogram (Figure 2c).

The biomarkers for resistance to allopurinol and antimonials determined by sequencing were verified by qPCR using LeishGenR^TM^, demonstrating high reliability (k > 0.8) and strong correlation for *metk* and *mrpa* copy numbers (*metk*: r = 0.914, p-value < 0.001; *mrpa*: r = 0.94, p-value < 0.001).

In parallel, virulence-associated CNVs were mainly conserved. Nine isolates (25%) showed a deletion in at least one gene of the mannose-containing glycocalyx biosynthesis pathway (*lpg1*, *ppm*, or *dpms*), which are essential for the complete synthesis of lipophosphoglycan (LPG) and proteophosphoglycan (PPG). Specifically, we observed a reduction to one copy of *lpg1* (*lpg1*+/-) in four out of 35 isolates (11%), *dpms* (*dpms*+/-) in one isolate (3%), and *ppm* (*ppm*+/-) in five isolates (14%). Among these, only CATB-102 exhibited a reduction in the copy number of two genes: *lpg1* (*lpg1*+/-) and *ppm* (*ppm*+/-), with no null mutants detected. All isolates retained both *lack* gene copies (*lack1*, *lack2*), which are implicated in virulence and immune response and are essential for vertebrate infection; therefore, the CNV (*lack*+-/--) associated with attenuation was not present in our cohort. However, isolates MCAN/IT/2020/1265 and MCAN/IT/2022/2885 exhibited an amplified *lack* genes copy number (5 copies).

biomarkers detected for each studied drug (allopurinol, amphotericin B, antimonial, miltefosine and paromomycin). b) 13 drug-resistance biomarkers profiles described in the cohort, 8 in HumL isolates (A-H) and 8 in CanL (A, B, E, I-M). Each profile is defined by a unique drug-resistance biomarkers status (presence/absence). Size is related to the prevalence of the drug resistance profiles. c) Heatmap showing the presence/absence of drug-resistance biomarkers (x-axis) within the *L. infantum* isolates identified in this study (y-axis). The presence of biomarkers is shown in orange and, absence in light yellow

### Locus-specific structure and repeat-driven rearrangements underpin the co-occurrence of drug resistance biomarkers in *L. infantum*

Hierarchical clustering of drug-resistance biomarkers revealed structured co-occurrence patterns, particularly among genes located within the same functional locus. The biomarkers for the extended H-locus and *METK* locus genes consistently clustered together, while *aqp1* and *LdMT* genes displayed independent variation (Figure 2c). To further understand the genomic basis of these CNV, we assessed whether local genomic rearrangements or broader chromosomal changes drove them. The analysis revealed three distinct correlation patterns when comparing gene copy numbers to chromosome somy. CNV at the extended H-locus (chr23) and *LdMT* (chr13) were strongly correlated with chromosomal somy (r = 0.88 and 0.76, respectively; p < 0.001), indicating that aneuploidy is a major driver of gene CN alteration. In contrast, *LdRos3*, *LdSMT*, *METK*, and *PMM* showed little or no correlation with somy, supporting a model of local structural rearrangement. Lastly, *AQP1* and *MSL*, located on tetrasomic chromosome 31, displayed moderate correlations, suggesting a complex interaction between chromosomal and sub-chromosomal events contributing to CNV (Additional file 3: Figure S1).

### Biomarker for antimonial resistance: amplification of the extended H-locus and reduction of the *aqp1* CNV

The extended H-locus biomarker, consisting of *abcc1*, *abcc2*, *mrpa*, and *ptr1*genes, was detected when the average gene copy number was >3.5, occurring in 9 of the 35 isolates.

Individually, *abcc1* and *mrpa* had >3.5 copies in nine isolates, *abcc 2* in eight, and *ptr1* in six. Notably, the nine isolates with the *mrpa* biomarker also had the extended H-locus biomarker (>3.5 copies) and >3 copies of chromosome 23, possibly indicating a sentinel marker for locus-wide amplification in cultures grown without drug stressors. These four biomarkers consistently co-occurred and formed a tight cluster in the heatmap analysis (Figure 2 c), suggesting co-amplification as a resistance mechanism.

In contrast, *aqp1* CNV reduction (<3.5) was detected in 13 isolates, clustering independently from the extended H-locus in the heatmap analysis (Figure 2 c), suggesting a mechanistically distinct resistance pathway. Among the 19 isolates positive for antimonial resistance biomarkers, the following profiles were detected: both extended H-locus and *aqp1* (4), extended H-locus only (5), *aqp1* only (8), *abcc1*, *mrpa*, and *aqp1* (1), *abcc2* (1) (Table 2).

To explore the structural basis of extended H-locus amplification, we analyzed the genomic context of chromosome 23 (coordinates 59,772–106,565, JPCM5 reference) (see Materials and Methods). Three repeated elements >0.5 kb were identified in the reference genome and in the *in silico* analysis of the 35 *de novo* assembled genomes. First, a 1.44 kb intergenic repeat sequence was identified three times (Rep01.01, Rep05.01, Rep06.01) with a nucleotide identity of greater than 99.5%. Rep01.01 was found to be an inverted repeat (IR), while Rep05.01 and Rep06.01 as direct repeats (DR) (Figure 3). Similar repeats have been reported previously [17, 18]. Second, a 0.5 kb sequence with 85% identity corresponding to an ATP-binding cassette domain, and third a 0.9 kb sequence with 90% identity across *abcc*1, *abcc2*, and *mrpa*, corresponding to an ABC transporter integral membrane type-1 and ATP binding cassette conserved domains (Figure 3).

**Figure 3.**
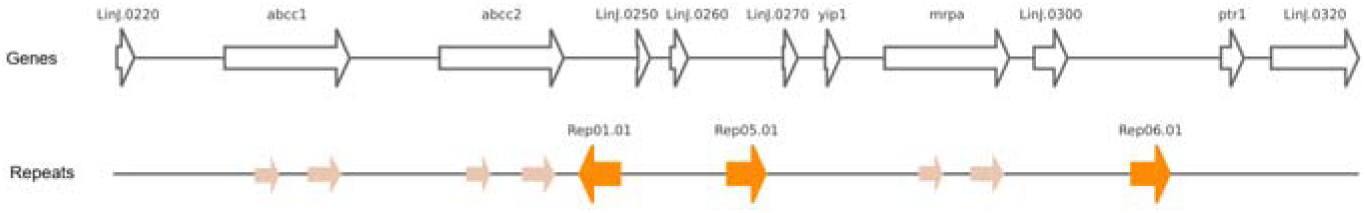
Extended H-locus structure in *L. infantum* and within the cohort. Genomic organization of the extended H-locus in *L. infantum*, with genes in white and intergenic repeats in orange (DR, Rep05.01 and Rep06.01; IR, Rep01.01). Homologous regions of 0.5 kb and 0.9 kb between genes are light orange.

Despite detecting these genome repeats, no potential extrachromosomal amplification elements from DR and IR were observed in our cohort ’s *de novo* assembled genomes, further supporting the notion that extended H-Locus CN correlates with chromosome 23 somy.

### Biomarker for allopurinol resistance: reduction of the *METK* Locus CNV

A total of 13 out of 35 isolates (37%) were positive for the allopurinol resistance biomarker, defined as having fewer than 1.5 copies of either *metk1* or *metk2*, encoding for a *S*-adenosylmethionine synthetase. All 13 were also positive for the *METK* locus resistance biomarker (<3 copies of *metk1* and *metk2*), reinforcing the reliability of this locus as a predictive biomarker. Notably, 85% (11/13) of these isolates were collected from dogs, highlighting a potential host-specific selection pressure. Among them, seven (63%) also tested positive for pentavalent antimonial biomarkers (Table 4). Hierarchical clustering supported this association, with *METK* and extended H-locus biomarkers often clustering together (Figure 2c). No isolates exhibited concurrent biomarkers for the combined treatment of allopurinol and miltefosine.

**Table 4.** Drug-resistance biomarkers to the combined allopurinol-antimonial treatment is the most frequent in allopurinol-resistant *L. infantum* CanL isolates. Summary of genomic resistance predisposition profiles to recommended combined drugs. Percentages are based on allopurinol-resistant isolates (n = 11).

| Genomic resistance predisposition | Resistance biomarker | Samples (%) |
| --- | --- | --- |
| Allopurinol + Antimonials | *METK* + extended H-locus/*aqp1* | 63 |
| Allopurinol + Miltefosine | *METK* locus + *LdMT/LdRos3/MSL* locus | 0 |
| Allopurinol only | *METK* locus | 37 |

The reduction in *metk* genes CNV results from a 5.1 kb deletion observed when mapping sequencing reads from the 13 samples to the JPCM5 reference genome. To understand the structural basis underlying the rearrangement of the *METK* locus, we conducted an *in silico* analysis of the JPCM5 reference sequence (RefSeq: GCF_000002875.2 ASM287v2), revealing the presence of two identical 5,120 bp repeat units (Rep.01.01, Rep.01.02), each encompassing a Lorien protein-coding gene (LinJ.30.3550/LinJ.30.3570) and a *metk* gene (LinJ.30.3560/LinJ.30.3580). Furthermore, the initial and final 218 bp of the 5,120 bp repeat are identical, indicating potential recombination hotspots (Figure 4). Read mapping revealed that positive isolates for the *METK* biomarker had lost one of the two repeat units. However, no precise breakpoint was defined, resulting in a hemizygous state on a disomic chromosome. Conversely, upon mapping the sequencing reads from the same isolates to their *de novo* assembled genomes, no evidence of deletion was observed, demonstrating continuous coverage and the absence of structural gaps. These findings suggest that the allopurinol resistance marker arises through a precise deletion event affecting one of the tandem 5,120 bp *METK* units, either the first or the second repeat unit on each chromosome.

**Figure 4.**
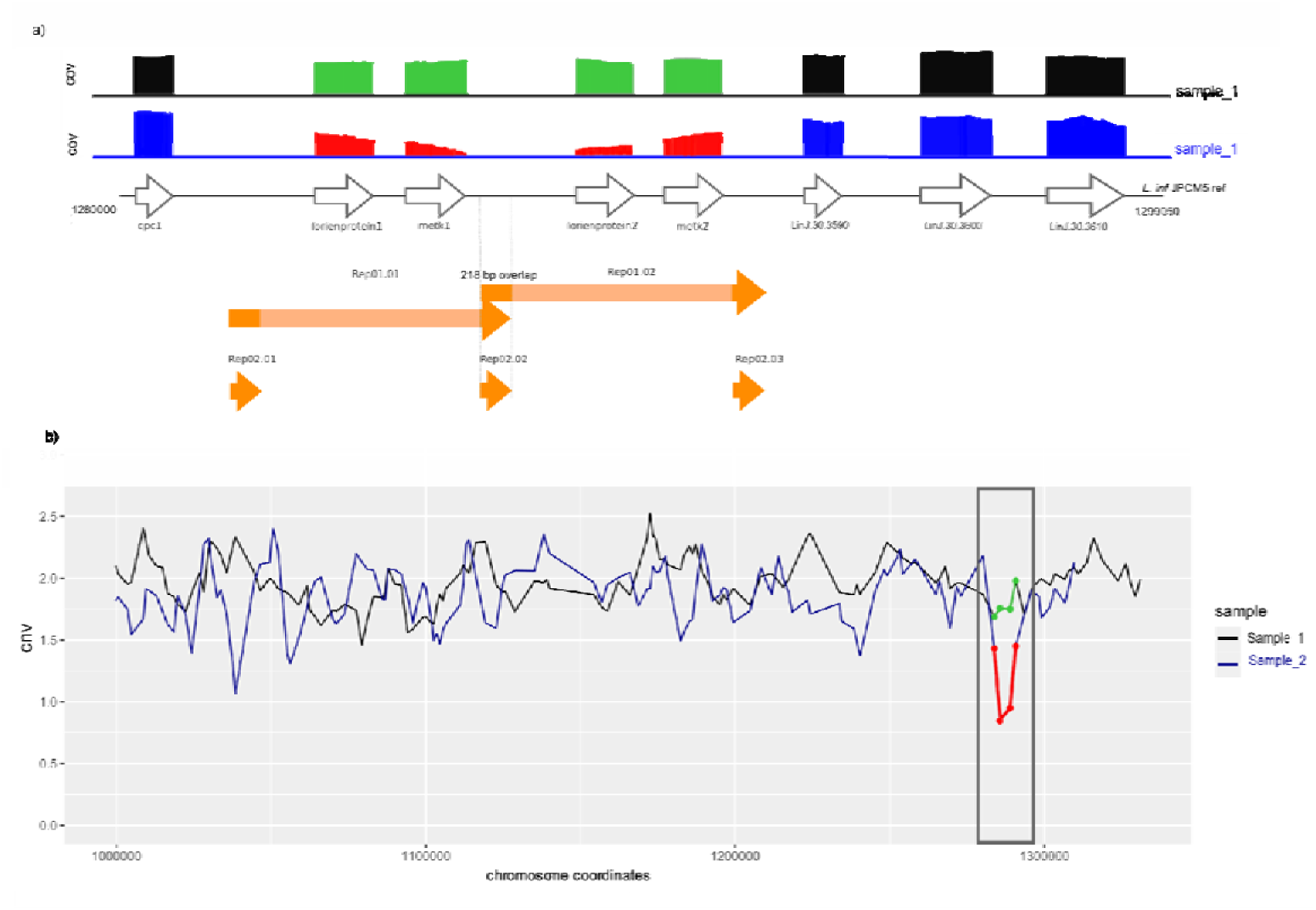
Extended H-locus structure in *L. infantum* and within the cohort. a) Characterization of the *METK* locus region of *L. infantum* and coverage decrease in the *METK* locus area due to 5.1 kb deletion, unlike the even coverage seen in samples with both constructs. B) CNV in *METK* locus.

### Biomarker for amphotericin B resistance: reduction of the *LdSMT* locus CNV

Genomic resistance biomarkers for amphotericin B are *LdSMT* genes, *smt1* and *smt2* at < 1.5 copies, encoding sterol 24-C-methyltransferases, and found in three (9%) isolates. Structural analysis and mapping of the sequencing reads of positive isolates to the SMT locus in the JPCM5 reference genome revealed that the reduction of *smt* genes CN was a result of a 3,789 bp deletion (3,786-3,792 bp) in the SMT locus mediated by homologous recombination between flanking repeats. The two near-identical (99.84% identity) *smt* genes were arranged in tandem and differ by a nucleotide at position 923 of the gene coordinate (*smt1*: (C); *smt2*: (T)). Each was encoded within a 1,268 bp repeat unit oriented in the same direction and including one *smt* gene, along with 26 nucleotides at the 5’ and 181 nucleotides at the 3’ end of the gene. A unique 2,521 bp intergenic sequence separated the two repeats, forming a total region of ∼3.8 kb (Figure 5).

**Figure 5.**
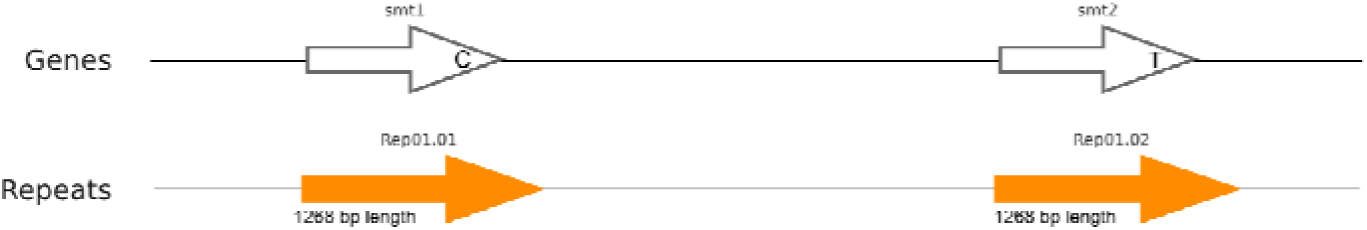
LdSMT locus structure in *L. infantum* and within the cohort. Genomic organization of the LdSMT locus.

Two positive isolates were from HumL, one of which (MHOM/TN/80/IPT-1) was positive for both biomarkers, while the other (MHOM/MA/67/ITMAP-264) tested positive only for the *smt1* gene. The third case, a canine isolate (MCAN/IL/2011/NT2), was near the positivity threshold for the *smt1* biomarker, with chromosome coverage approaching the quality inclusion criteria for LeishGenApp (1.33 copies, chr. cov = 16X). Sequence alignment of deletion-containing reads of the three isolates revealed highest identity with *smt2* (XM_001469794.1), which presented a thymine (T) at position 923, confirming that *smt1* (cytosine at position 923) was the gene lost.

### Drug-resistance biomarkers to miltefosine: *LdMT* and *LdRos3* genes

Drug resistance biomarkers to miltefosine were detected in three of the 35 *L. infantum* isolates through CNV detected in the *LdMT* (two isolates with <1.5 copies) and the *LdRos3* (one isolate with <1.5 copies) genes. *LdMT* encodes for a P-type ATPase phospholipid translocase, while *LdRos3* encodes for the β subunit of LdMT. Both proteins are part of the same flippase machinery, and not individually sufficient for the translocation of miltefosine (Pérez-Victoria et al., 2006). Therefore, these three samples would be similarly resistant to miltefosine due to a decrease in drug accumulation, as they are defective in the uptake of miltefosine by being deficient in one of the required *LdMT* and *LdRos3* (Pérez-Victoria et al., 2006).

### Drug-resistance biomarkers to paromomycin: *b-cat* and *d-ldh* genes

The drug resistance biomarker for paromomycin was determined as <1.5 copies of *b-cat,* and <1.5 copies of *d-ldh* (Table 2). Eight of 35 samples were positive for at least one resistance biomarker, and four had both biomarkers. Across the positive isolates, 62,5% (5/8) were from HumL, while 37,5% from CanL. Significantly, three positive HumL isolates were also positive for antimonial, present in profiles C and D.

## Discussion

### Detection of aneuploidies and genome plasticity in *L. infantum* genome by nanopore-only whole-genome sequencing

Whole-genome sequencing with long-read nanopore technology revealed pervasive aneuploidy and widespread mosaicism across the 35 *L. infantum* isolates, providing sufficient resolution to reconstruct karyotypes. The diverse range of aneuploidies observed in this study is consistent with previous studies using next-generation sequencing data from *Leishmania* spp., which report frequent aneuploidies in this species [64, 65], even without drug pressure [41]. Only one isolate exhibited the canonical disomic karyotype with tetrasomy of chromosome 31, emphasizing the degree of genomic plasticity within the field population. Not all chromosomes hold the same aneuploidy turnover potential. A subset of chromosomes (7, 14, 18, 19, 27, 28, 34, and 36) remained disomic and stable, while chromosomes 8 and 23 were potentially aneuploid. The results are consistent across five of the stable chromosomes (19, 27, 28, 34, and 36), as well as in both chromosomes (8 and 23), which exhibit significant CNV when compared with a study involving 161 cultured samples. [64]. Trisomy of chromosomes 8, 20, and 23, and tetrasomy of chromosome 31, were frequent across the dataset, consistent with previous reports [64, 66]. In agreement with those studies, monosomy was rare and often transient. Our analysis further supports the model of mosaicism, where non-integer somy values reflect a population-level mixture of karyotypes within the same isolate, which is important in stress adaptation and natural selection [66].

The present data suggest that mid-term culturing (up to 33 passages) does not have a significant effect on aneuploidy levels compared to cultured clinical isolates (< 10 passages), with culturing beyond 30 passages potentially affecting chromosomal plasticity, as seen in long-established clonal lines MHOM/TN/80/IPT-1 and MCAN/IT/2020/1265 with 10 and 11 supernumerary chromosomes, respectively. This is beneficial for clinicians, suggesting that *in vitro* isolates resemble the *Leishmania* population in the infected host. For instance, aneuploidy in chromosome 23 is one of the most common variations observed in field isolates, and one of the rapidly established trisomy *in vitro* [66]. Longitudinal studies monitoring clinical isolates and *in vitro* aneuploidy dynamics are of interest [66]. Assessing the potential effect of different culture media would also be valuable in clarifying the thresholds at which culturing may impact *Leishmania*.

As aneuploidy could contribute to parasite resistance, monitoring of karyotype patterns could inform treatment protocols. In the context of antimonial resistance, it is associated with aneuploidy of chromosomes 1, 11, and 25 [17]. The authors reported an increase in the total DNA content of the chromosomes, which correlates with the mRNA profile and gene overexpression. the cultures MHOM/MA/67/ITMAP-264 and MHOM/MA/67/ITMAP-265 exhibited polyploid characteristics in chromosomes 1 and 11. Both cultures derived from the same starting population, although the latter underwent 10 more passages. Gene CNV, identified as biomarkers for antimony resistance, were present in MHOM/MA/67/ITMAP-264 (*abcc1, abcc2, mrpa*) and MHOM/MA/67/ITMAP-265 (*abcc1, abcc2, mrpa, aqp1*). Conversely, no aneuploidy for chromosomes 1, 11, or 25 was observed in other cultures presenting CNV. According to collaborators [17], the withdrawal of drug pressure resulted in the reversion of supernumerary chromosomes 1, 11, and 25 to their original somy levels. This observation suggests that these chromosomal adaptations were not actively developed in our isolates, as our cultures were not subjected to drug pressure during the culturing process.

### *L. infantum* isolates from dogs and humans show distinct drug-resistance biomarker status while mostly preserving virulence

In the current study, 80% (28/35) of *L. infantum* isolates harbored at least one drug resistance-associated genomic biomarker, with non-negligible values of genomic resistance biomarker prevalence in *L. infantum* isolates across the Mediterranean. It is worth considering that the drug resistance biomarkers detected in our study rely on *in silico* prediction, without experimental validation (i.e., inhibitory concentrations, IC50) for all the samples tested, which may represent a limitation of our study. The observed prevalences could not be compared with existing data due to the lack of published literature assessing the drug-resistance status of *L. infantum* strains circulating in the Mediterranean.

When considering the prevalence of drug resistance biomarkers, it is noteworthy to consider sample size limitations and potential geographic representation bias. Additionally, the type of culture media used by different institutions could also interfere with *Leishmania* CNV, representing a source of bias in the analysis. Culture-independent approaches for biomarker detection direct from clinical samples, such as gene CNV detection by Real-Time PCR or sequencing, should be of interest for further studies to avoid culture adaptation changes and culture media effects on CNV associated with Leishmania genome plasticity.

The observation that there were differences in genomic resistance biomarkers among the five studied drugs was of interest, as it underscores the importance of drug pressure, sandfly transmission implications, and targeted drug policies in mitigating the dissemination of resistance. Antimonial resistance biomarkers appeared to be the highest in both canine (50%) and human (61%) *L. infantum* isolates. The most likely explanation for this is the earlier use of pentavalent antimonials for CanL and HumL treatment [5], the retention of the resistant phenotype through sand flies, and demonstrated enhanced fitness compared to sensitive strains [67] This scenario places antimonial-resistant strains as a substantial threat in areas where antimony drug pressure is sought to be low and underscores the need for careful regulation of antimonial use to control the spread of resistance. Particularly in dogs, highlighting their role as primary reservoirs, lessening clinical relapses due to resistance, and their implications in sand fly transmission. Contrary, miltefosine resistance was notably low (9%), likely due to two factors: i) miltefosine-resistant parasites exhibit reduced fitness in mammalian hosts, limiting their spread despite maintaining their phenotype through the vector [68], and ii) miltefosine is not used in HumL treatment, reducing selective pressure [5]. These findings support the WHO recommendations to restrict miltefosine use to CanL, thereby preventing the emergence of resistance in humans while preserving its efficacy in dogs. For paromomycin, the observed resistance prevalence was higher in humans (38%) than in dogs (14%). Resistant parasites maintain their phenotype through sand flies [69], but reduced fitness in the absence of drug pressure may explain their limited dissemination in CanL.

The pairwise comparison reported herein for drug-resistance positivity between dogs and humans revealed that host-specific patterns seem to emerge for the resistance profiles (P = 0.08), and the allopurinol resistance biomarker (P = 0.07), despite no statistically significant differences being found. Our study would have benefited from an increase in sample size and a balanced number of samples from each host group. Additional studies with an increased sample size will enable confirm the insights extracted from our studied cohort. These marked differences between host groups potentially reflect therapeutic practice and host-specific drug exposure while partially explaining the clinical performance of each *L. infantum* strain.

Scanning the drug resistance biomarkers in canine isolates revealed that allopurinol and antimonials were the most common, each at a frequency of 50%. Allopurinol is the primary medication for the long-term management of CanL cases [70]. Notably, there is evidence suggesting that allopurinol exerts a pressure effect as a driver of *METK* locus CNV. *METK* positivity in HumL isolates is significantly lower at 15%, probably due to the limited use of allopurinol for HumL management [5].

This study aimed to investigate the potential link between the recommended combined therapy for CanL, which includes the leishmaniostatic drug allopurinol and either meglumine antimoniate or miltefosine as leishmanicidal drugs, and the natural emergence of genomic resistance in field isolates. Notably, the current data showed that among isolates with allopurinol resistance biomarkers, a high proportion (63%) also possessed genomic resistance biomarkers to the combined therapy of antimonial and allopurinol, aligning well with the earliest use of these drugs [71]. On the other hand, our results showed no isolates showing genomic resistance biomarkers to the combined therapy of allopurinol and miltefosine, supporting the continued efficacy of this combination therapy in CanL. We detected only one *in vitro* clinical isolate with biomarkers of miltefosine resistance (MCAN/IT/2019/3299). Sample 3299 was isolated in 2019 after the acceptance of miltefosine for CanL treatment. Knowing whether this canine subject had been treated with miltefosine before sample collection would be valuable as miltefosine has a long half-life in the treated host, possibly leading to the retention of sub-therapeutic levels in the body (Pérez-Victoria et al., 2006), which may trigger the development of resistance [15]. The prevalence of low miltefosine resistance biomarkers aligned with the later approval of miltefosine for CanL treatment in Europe (2007) and the few existing clinically resistant cases in the literature, the first reported in 2020 [15].

Regarding HumL, genomic resistance biomarkers to antimonials were the most common, followed by resistance to paramomycin. These findings align with current treatment guidelines for both visceral (VL) and cutaneous leishmaniasis (CL). Pentavalent antimonials and liposomal amphotericin B are the primary treatments for VL. Concerning CL caused by L. infantum in the Mediterranean basin, paromomycin ointment, despite its late approval in 2006, and intralesional antimonials are used as topical treatments, while pentavalent antimonials continue to be the first-line systemic option [5]. Notably, our research revealed an interestingly low prevalence of genomic resistance biomarkers to amphotericin B (15%). This underscores the importance of the WHO recommendations for European veterinarians to avoid using this drug, particularly as efficacy is not completely clear in dogs [72], and they may develop higher fitness-resistant parasites [73] that can be transmitted to humans.

Hierarchical clustering uncovered potentially different drug-resistance profiles between hosts. Dogs predominantly exhibited monoresistance (50%), especially to allopurinol, while resistance to two drugs was more frequently detected in human isolates (38%). Treatment guidelines for CanL recommend combined therapy with at least two drugs with different mechanisms of action to delay drug resistance [74, 75], while for HumL, monotherapy is the recommended treatment approach [5]. Our findings support the effectiveness of the veterinarian’s strategy. However, resistance to combination therapy is also possible, with 31% of canine isolates showing genomic resistance to two or three drugs, consistent with previous reports [76]. Differences with HumL isolates suggest a potential influence of therapeutic strategies on the emergence of resistance. However, since a significant portion of two-drug resistant strains (80%) had antimonial resistance biomarkers, it suggests preadaptation, possibly caused by historical drug exposure in old-world *Leishmania* circulating in humans [46, 77]. This potential preadaptation could act as a confounding factor in assessing the effect of treatment guidelines on the development of drug resistance. Therefore, further clinical trials on combination therapy for HumL in the Mediterranean region are necessary to clarify how therapeutic strategies influence resistance profiles.

A limited number of attenuation and virulence biomarkers were identified in the studied isolates, even in the presence of drug-resistance biomarkers. A reduction of one copy in *ppm*, *dpms*, and *lpg1* was observed in some of the isolates. The *ppm* gene encodes phosphomannomutase, an enzyme essential for GDP-Man synthesis and indirectly involved in LPG and PPG biosynthesis[21]. The *dpms* gene encodes Dol-P-Man synthase, while the *lpg1* gene encodes a galactofuranosyltransferase; the former supplies the β-Man donor necessary for initial LPG mannose [21], and the latter adds side chains essential for mature LPG structure [22, 55, 58]. Null mutants of these genes lack or produce truncated PPG and LPG molecules [21]. Investigating the potential impact of a single-copy reduction on the disruption of side-chain formation and subsequent alterations in glycocalyx surface composition will be of particular interest, as these changes may influence sand fly midgut adhesion, macrophage recognition, or immune evasion. Notably, all isolates preserved two intact copies of *lack1* and *lack2*, supporting previous evidence that Lack protein is essential for intracellular survival and infectivity in mammalian hosts [59–61, 63]. Further studies to understand the role of increased *lack* genes copy number in parasite virulence and its potential link to higher parasite loads and clinical stage will be of particular interest.

### CNV for drug resistance biomarkers in *L. infantum*

The present study also investigated whether exclusively local genomic rearrangements drift the CNV in key drug target and transporter genes or if they are influenced by the already discussed aneuploidy, detecting three different correlation patterns.

Of interest, the chromosome 23 somy showed a strong positive correlation with the copy number of the extended H-locus genes, suggesting that aneuploidy events are drivers of these genes’ CNV, with nine isolates having >3 copies of both chromosome 23 and extended H-locus. Although described as a highly unstable chromosome, *L. infantum* isolates with up to 3 copies of chromosome 23 are known to be sensitive to antimonials without extrachromosomal amplifications [46]. Based on the evidence from the literature, the genomic resistance biomarkers threshold was set as >3.5 copies for the four extended H-locus genes in chromosome 23, contrary to other biomarkers encoded in disomic chromosomes (see Table 1). Chromosome 23 consistently displayed an increase in copy number in *in vitro* populations selected for antimonial resistance in which *mrpa* was artificially deleted [47] suggesting an adaptive advantage as it is known as a rapid way to amplify a cassette of genes involved in the antimonials pathway. Our results pointed out that, beyond the intrinsic variability of the chromosome, the widespread increase in the dosage of chromosome 23 observed in the cohort may represent a form of pre-adaptation to trivalent antimonials, likely driven by the sustained antimonial pressure exerted overuse decades in the Mediterranean basin [77].

Structural analysis of the extended H-locus revealed the presence of a sequence repeated three times (Rep01.01, Rep05.01, and Rep06.01) in the intergenic region, consistent with previous studies [17, 18]. Also, the presence of two sequences repeated twice in the coding regions. It is well described that widely distributed repeated sequences, mainly in non-coding regions of the *Leishmania* genome, lead to stochastic genome rearrangements [18, 66] playing a crucial role in gene amplification or expansion and gene deletion, determining parasite’s adaptability to drug stress. The three repeated sequences described in the extended H-locus suggest the potential formation of 4 circular episomes and two linear minichromosomes, depending on the stochastic rearrangement. However, no episomal amplification was detected in our cohort. The promastigote cultures analyzed were grown under natural conditions. In contrast, experimental episomal amplification has mainly been detected in cultures grown under drug pressure and demonstrated that it generally disappears rapidly when the drug pressure is removed [17] [18] [78] [79]. On the other hand, we cannot discard the effect of the DNA extraction procedure used as a drawback to episomal recovery.

Concerning allopurinol resistance, a 5.12 kb repeated sequence flanked by 218 bp repeats was detected on chromosome 30 in the *METK* locus. This 218 bp intergenic repeated sequence may have been a determinant in the rearrangement of the *L. infantum METK* locus structure, as observed in the reference genome and 21 samples (containing two identical 5.12 kb copies in tandem). That 218 bp repeat might be sufficient in length for the loss of a 5.12 kb *METK* region by homologous recombination, similar to the 220bp direct repeat at the start codon of the tartrate-sensitive acid phosphatase (*LdBPK_366740*) and histidine secretory acid phosphatase (*LdBPK_366770*) that enables the MAPK locus gene amplification in antimonial resistant lines, maintained as a 1,700bp episomal DNA [80]. Moreover, a minimum identity of 85% is required for recombination in *Leishmania* [81], and our results showed a 97-100% identity. Therefore, the *METK* and the *SMT* locus studied are examples of gene amplification strategies, as two identical copies are in tandem.

Leveraging WGS, our approach provides a methodological advantage by complementing CNV analysis with detailed read sequence profiling, with significant implications for the SMT locus. We confirmed *smt1* loss and distinguished it from *smt2*, which differs by a single nucleotide. This finding may have drug-resistance implications, as a previous study described *smt1* loss as the cause of amphotericin B resistance, with *smt2* potentially compensating [51].

## Conclusion

Detecting resistance biomarkers in *L. infantum* could improve of patient management by enabling timely and precise therapy, identifying resistant parasites and subpopulations in CanL and HumL. This allows case selection based on genomic resistance, potentially reducing the risk of relapses and the spread of resistance and thereby prolonging the therapeutic lifespan of current drugs. Resistant *L. infantum* strains in canines, the primary host and main reservoir, hinder the control of these strains by facilitating transmission to dogs and humans. Expanding the spectrum of gene CNV described as genomic resistance and pathogenicity biomarkers, combined with the detection of single-point mutations associated with drug resistance, can improve LeishGenApp’s accuracy, aiding therapy decisions. In addition, conducting molecular *in vitro* analysis (i.e., inhibitory concentrations, IC50) of the studied cultures may be of interest to establish a robust correlation between the genomic resistance biomarkers identified through WGS and the phenotypic traits.

## Supplementary information

**Additional file 1: Table S1.** N50, total genome size, GC content, and the number of contigs for each *de novo* assembly. **Additional file 2: Table S2.** CNV and drug resistance biomarker detection. **Additional file 2: Figure S1.** Correlation between chromosome copy number and biomarker copy number (sub-chromosomal level).

## Abbreviations

CNV: Copy Number Variation; CN: Copy Number; *L. infantum: Leishmania infantum*; WGS: Whole-Genome Sequencing; HumL: Human leishmaniosis; CanL: Canine leishmaniosis; DR: Direct repeat: IR: inverted repeat.

## Declarations

### Acknowledgments

The authors would like to thank Lorena Bernardo and Jose Carlos Solana from Instituto de Salud Carlos III, Ana Isabel Pinto from Instituto de Investigação e Inovação em Saúde (i3S), as well as Germano Castelli from Istituto Zooprofilattico sperimentale della Sicilia for their contributions in preparing the *L. infantum* strains in the laboratory.

The authors would like to thank the anonymous reviewers for their comments and suggestions, which have improved the reporting of our results.

### Availability of data and material

The datasets supporting the findings of this article are included within the article, and the sequencing data are available on the DDBJ/ENA/GenBank repository under BioProject PRJNA881045, with the following accession numbers: JAZFNU000000000-JAZFNZ000000000, JAZFOA000000000-JAZFOZ000000000, and JAZFPA000000000-JAZFPC000000000.

### Author contributions

MCM, JMC, OF, XR and LF conceived and designed the experiments. MG, MA, GB, FB, CC, JC, TD, CM, AP, XRG, NS, AVS, and DYL assisted with the collection and sending of *L. infantum* isolates. MCM and OF conducted the formal analysis. MCM carried out the sequencing experiments, collected the data required for the study and performed the analysis. MCM and OF wrote the original draft. OF, XR, JMC, LF, GB, AC, TD, CM, JM and, FV reviewed the manuscript. All authors read and approved the final version of the manuscript.

### Ethics approval and consent to participate

Not applicable.

### Consent for publication

Not applicable.

## Bibliography

1. Ponte-Sucre A, Gamarro F, Dujardin JC, Barrett MP, López-Vélez R, García-Hernández R, et al. Drug resistance and treatment failure in leishmaniasis: A 21st century challenge. PLoS Neglected Tropical Diseases. 2017;11.

2. Maia C, Conceição C, Pereira A, Rocha R, Ortuño M, Muñozid C, et al. The estimated distribution of autochthonous leishmaniasis by *Leishmania infantum* in Europe in 2005–2020. PLoS Negl Trop Dis. 2023;17.

3. Moreno J, Alvar J. Canine leishmaniasis: epidemiological risk and the experimental model. Trends Parasitol. 2002;18:399–405.

4. Gupta AK, Das S, Kamran M, Ejazi SA, Ali N. The pathogenicity and virulence of *Leishmania* - interplay of virulence factors with host defenses. Virulence. 2022;13.

5. Gradoni L, Lopez-Vélez R, Mokni M. Manual on case management and surveillance of the leishmaniases in the who european region. 2017.

6. Pérez-Victoria FJ, Sánchez-Cañete MP, Seifert K, Croft SL, Sundar S, Castanys S, et al. Mechanisms of experimental resistance of *Leishmania* to miltefosine: Implications for clinical use. Drug Resistance Updates. 2006;9:26–39.

7. Matos AP, Viçosa AL, Ré MI, Ricci-Júnior E, Holandino C. A review of current treatments strategies based on paromomycin for leishmaniasis. Journal of Drug Delivery Science and Technology. 2020;57.

8. Kaempfle M, Hartmann K, Bergmann M. Treatment of *Leishmania infantum* Infections in Dogs. Microorganisms. 2025;13.

9. Ibarra-Meneses AV, Corbeil A, Wagner V, Beaudry F, do Monte-Neto RL, Fernandez-Prada C. Exploring direct and indirect targets of current antileishmanial drugs using a novel thermal proteomics profiling approach. Front Cell Infect Microbiol. 2022;12.

10. Jeddi F, Mary C, Aoun K, Harrat Z, Bouratbine A, Faraut F, et al. Heterogeneity of molecular resistance patterns in antimony-resistant field isolates of l*eishmania* species from the western mediterranean area. Antimicrob Agents Chemother. 2014;58.

11. Leprohon P, Fernandez-Prada C, Gazanion É, Monte-Neto R, Ouellette M. Drug resistance analysis by next generation sequencing in *Leishmania*. International Journal for Parasitology: Drugs and Drug Resistance. 2015;5.

12. Gramiccia M, Gradoni L, Orsini S. Decreased sensitivity to meglumine antimoniate (Glucantime) of *Leishmania infantum* isolated from dogs after several courses of drug treatment. Ann Trop Med Parasitol. 1992;86.

13. Yasur-Landau D, Jaffe CL, David L, Baneth G. Allopurinol Resistance in *Leishmania infantum* from Dogs with Disease Relapse. PLoS Negl Trop Dis. 2016;10.

14. Schäfer I, Faucher M, Nachum-Biala Y, Ferrer L, Carrasco M, Kehl A, et al. Evidence for in vivo resistance against allopurinol in a dog infected with *Leishmania infantum* by reduction in copy numbers of the S-adenosylmethionine synthetase (METK) gene. Parasit Vectors. 2024;17.

15. Gonçalves G, Campos MP, Gonçalves AS, Medeiros LCS, Figueiredo FB. Increased *Leishmania infantum* resistance to miltefosine and amphotericin B after treatment of a dog with miltefosine and allopurinol. Parasit Vectors. 2021;14.

16. Cojean S, Houzé S, Haouchine D, Huteau F, Lariven S, Hubert V, et al. *Leishmania resistance* to miltefosine associated with genetic marker. Emerging Infectious Diseases. 2012;18.

17. Leprohon P, Légaré D, Raymond F, Madore É, Hardiman G, Corbeil J, et al. Gene expression modulation is associated with gene amplification, supernumerary chromosomes and chromosome loss in antimony-resistant *Leishmania infantum*. Nucleic Acids Res. 2009;37.

18. Ubeda JM, Raymond F, Mukherjee A, Plourde M, Gingras H, Roy G, et al. Genome-Wide Stochastic Adaptive DNA Amplification at Direct and Inverted DNA Repeats in the Parasite *Leishmania*. PLoS Biol. 2014;12.

19. Pérez-Victoria FJ, Gamarro F, Ouellette M, Castanys S. Functional cloning of the miltefosine transporter: A novel p-type phospholipid translocase from leishmania involved in drug resistance. Journal of Biological Chemistry. 2003;278.

20. Yasur-Landau D, Jaffe CL, David L, Doron-Faigenboim A, Baneth G. Resistance of *Leishmania infantum* to allopurinol is associated with chromosome and gene copy number variations including decrease in the S-adenosylmethionine synthetase (METK) gene copy number. Int J Parasitol Drugs Drug Resist. 2018;8.

21. Garami A, Mehlert A, Ilg T. Glycosylation Defects and Virulence Phenotypes of Leishmania mexicana Phosphomannomutase and Dolicholphosphate-Mannose Synthase Gene Deletion Mutants . Mol Cell Biol. 2001;21.

22. Späth GF, Epstein L, Leader B, Singer SM, Avila HA, Turco SJ, et al. Lipophosphoglycan is a virulence factor distinct from related glycoconjugates in the protozoan parasite *Leishmania major*. Proc Natl Acad Sci U S A. 2000;97.

23. Ilgoutz SC, McConville MJ. Function and assembly of the *Leishmania* surface coat. In: International Journal for Parasitology. 2001.

24. Azevedo LG, De Queiroz ATL, Barral A, Santos LA, Ramos PIP. Proteins involved in the biosynthesis of lipophosphoglycan in *Leishmania*: A comparative genomic and evolutionary analysis. Parasit Vectors. 2020;13.

25. Martí-Carreras J, Carrasco M, Gómez-Ponce M, Noguera-Julián M, Fisa R, Riera C, et al. Identification of *Leishmania infantum* Epidemiology, Drug Resistance and Pathogenicity Biomarkers with Nanopore Sequencing. Microorganisms. 2022;10.

26. González-De La Fuente S, Peiró-Pastor R, Rastrojo A, Moreno J, Carrasco-Ramiro F, Requena JM, et al. Resequencing of the *Leishmania infantum* (strain JPCM5) genome and *de novo* assembly into 36 contigs. Sci Rep. 2017;7.

27. Ren J, Chaisson MJP. lra: A long read aligner for sequences and contigs. PLoS Comput Biol. 2021;17.

28. Li H, Handsaker B, Wysoker A, Fennell T, Ruan J, Homer N, et al. The Sequence Alignment/Map format and SAMtools. Bioinformatics. 2009;25.

29. Talevich E, Shain AH, Botton T, Bastian BC. CNVkit: Genome-Wide Copy Number Detection and Visualization from Targeted DNA Sequencing. PLoS Comput Biol. 2016;12.

30. Patino LH, Imamura H, Cruz-Saavedra L, Pavia P, Muskus C, Méndez C, et al. Major changes in chromosomal somy, gene expression and gene dosage driven by SbIII in *Leishmania braziliensis* and *Leishmania panamensis*. Sci Rep. 2019;9.

31. Choudhury A, Hazelhurst S, Meintjes A, Achinike-Oduaran O, Aron S, Gamieldien J, et al. Population-specific common SNPs reflect demographic histories and highlight regions of genomic plasticity with functional relevance. BMC Genomics. 2014;15.

32. De Coster W, Rademakers R. NanoPack2: population-scale evaluation of long-read sequencing data. Bioinformatics. 2023;39.

33. Kolmogorov M, Yuan J, Lin Y, Pevzner PA. Assembly of long, error-prone reads using repeat graphs. Nat Biotechnol. 2019;37.

34. Manni M, Berkeley MR, Seppey M, Zdobnov EM. BUSCO: Assessing Genomic Data Quality and Beyond. Curr Protoc. 2021;1.

35. Steinbiss S, Silva-Franco F, Brunk B, Foth B, Hertz-Fowler C, Berriman M, et al. Companion: a web server for annotation and analysis of parasite genomes. Nucleic Acids Res. 2016;44.

36. Marçais G, Delcher AL, Phillippy AM, Coston R, Salzberg SL, Zimin A. MUMmer4: A fast and versatile genome alignment system. PLoS Comput Biol. 2018;14.

37. Camacho C, Coulouris G, Avagyan V, Ma N, Papadopoulos J, Bealer K, et al. BLAST+: Architecture and applications. BMC Bioinformatics. 2009;10.

38. Thorvaldsdóttir H, Robinson JT, Mesirov JP. Integrative Genomics Viewer (IGV): High-performance genomics data visualization and exploration. Brief Bioinform. 2013;14.

39. Wickham H, Averick M, Bryan J, Chang W, McGowan L, François R, et al. Welcome to the Tidyverse. J Open Source Softw. 2019;4.

40. Cui Y, Chen X, Luo H, Fan Z, Luo J, He S, et al. BioCircos.js: An interactive Circos JavaScript library for biological data visualization on web applications. Bioinformatics. 2016;32.

41. Rogers MB, Hilley JD, Dickens NJ, Wilkes J, Bates PA, Depledge DP, et al. Chromosome and gene copy number variation allow major structural change between species and strains of Leishmania. Genome Res. 2011;21.

42. Pérez-Victoria FJ, Sánchez-Cañete MP, Castanys S, Gamarro F. Phospholipid translocation and miltefosine potency require both *L. donovani* miltefosine transporter and the new protein LdRos3 in Leishmania parasites. Journal of Biological Chemistry. 2006;281.

43. Carnielli JBT, Crouch K, Forrester S, Silva VC, Carvalho SFG, Damasceno JD, et al. A *Leishmania infantum* genetic marker associated with miltefosine treatment failure for visceral leishmaniasis. EBioMedicine. 2018;36:83–91.

44. Douanne N, Wagner V, Roy G, Leprohon P, Ouellette M, Fernandez-Prada C. MRPA-independent mechanisms of antimony resistance in *Leishmania infantum*. Int J Parasitol Drugs Drug Resist. 2020;13:28–37.

45. El Fadili K, Messier N, Leprohon P, Roy G, Guimond C, Trudel N, et al. Role of the ABC transporter MRPA (PGPA) in antimony resistance in *Leishmania infantum* axenic and intracellular amastigotes. Antimicrob Agents Chemother. 2005;49.

46. Dumetz F, Cuypers B, Imamura H, Zander D, D’Haenens E, Maes I, et al. Molecular Preadaptation to Antimony Resistance in *Leishmania donovani* on the Indian Subcontinent. mSphere. 2018;3.

47. Negreira GH, de Groote R, Van Giel D, Monsieurs P, Maes I, de Muylder G, et al. The adaptive roles of aneuploidy and polyclonality in *Leishmania* in response to environmental stress . EMBO Rep. 2023;24.

48. Gourbal B, Sonuc N, Bhattacharjee H, Legare D, Sundar S, Ouellette M, et al. Drug uptake and modulation of drug resistance in *Leishmania* by an aquaglyceroporin. Journal of Biological Chemistry. 2004;279.

49. Andrade JM, Baba EH, Machado-De-Avila RA, Chavez-Olortegui C, Demicheli CP, Frézard F, et al. Silver and nitrate oppositely modulate antimony susceptibility through aquaglyceroporin 1 in *Leishmania (Viannia*) species. Antimicrob Agents Chemother. 2016;60:22–60.

50. Rastrojo A, García-Hernández R, Vargas P, Camacho E, Corvo L, Imamura H, et al. Genomic and transcriptomic alterations in *Leishmania donovani* lines experimentally resistant to antileishmanial drugs. Int J Parasitol Drugs Drug Resist. 2018;8.

51. Tulloch LB, Tinti M, Wall RJ, Weidt SK, Corpas- Lopez V, Dey G, et al. Sterol 14-alpha demethylase (CYP51) activity in *Leishmania donovani* is likely dependent upon cytochrome P450 reductase 1. PLoS Pathog. 2024;20:e1012382.

52. Jesus-Santos FH, Lobo-Silva J, Ramos PIP, Descoteaux A, Lima JB, Borges VM, et al. LPG2 Gene Duplication in *Leishmania infantum*: A Case for CRISPR-Cas9 Gene Editing. Front Cell Infect Microbiol. 2020;10.

53. Azevedo LG, De Queiroz ATL, Barral A, Santos LA, Ramos PIP. Proteins involved in the biosynthesis of lipophosphoglycan in *Leishmania*: A comparative genomic and evolutionary analysis. Parasit Vectors. 2020;13.

54. Garami A, Mehlert A, Ilg T. Glycosylation Defects and Virulence Phenotypes *of Leishmania mexicana* Phosphomannomutase and Dolicholphosphate-Mannose Synthase Gene Deletion Mutants . Mol Cell Biol. 2001;21:8168–83.

55. Forestier CL, Gao Q, Boons GJ. Leishmania lipophosphoglycan: How to establish structure-activity relationships for this highly complex and multifunctional glycoconjugate? Front Cell Infect Microbiol. 2014;4.

56. Ilgoutz SC, Mcconville MJ. Invited review Function and assembly of the *Leishmania* surface coat.

57. Spä GF, Epstein L, Leader B, Singer SM, Avila HA, Turco SJ, et al. Lipophosphoglycan is a virulence factor distinct from related glycoconjugates in the protozoan parasite Leishmania major. 2000.

58. Lázaro-Souza M, Matte C, Lima JB, Duque GA, Quintela-Carvalho G, Vivarini Á de C, et al. *Leishmania infantum* lipophosphoglycan-deficient mutants: A tool to study host cell-parasite interplay. Front Microbiol. 2018;9 APR.

59. Sánchez-Sampedro L, Gómez CE, Mejías-Pérez E, Sorzano CO, Esteban M. High quality long-term CD4+ and CD8+ effector memory populations stimulated by DNA-LACK/MVA-LACK regimen in *Leishmania major* BALB/C model of infection. PLoS One. 2012;7.

60. Sjölander A, Baldwin TM, Curtis JM, Handman E. Induction of a Th1 Immune Response and Simultaneous Lack of Activation of a Th2 Response Are Required for Generation of Immunity to Leishmaniasis 1. 1998.

61. Kelly BL, Stetson DB, Locksley RM. Leishmania major LACK Antigen Is Required for Efficient Vertebrate Parasitization. Journal of Experimental Medicine. 2003;198:1689–98.

62. Jensen KDC, Sercarz EE, Gabaglia CR. Altered peptide ligands can modify the Th2 T cell response to the immunodominant 161-175 peptide of LACK (*Leishmania* homolog for the receptor of activated C kinase). Mol Immunol. 2009;46:366–74.

63. Julia V, Rassoulzadegan M, Glaichenhaus N. Resistance to *Leishmania major* Induced by Tolerance to a Single Antigen. 1996.

64. Franssen SU, Durrant C, Stark O, Moser B, Downing T, Imamura H, et al. Global genome diversity of the *leishmania donovani* complex. Elife. 2020;9.

65. Negreira GH, Monsieurs P, Imamura H, Maes I, Kuk N, Yagoubat A, et al. High throughput single-cell genome sequencing gives insights into the generation and evolution of mosaic aneuploidy in *Leishmania donovani*. Nucleic Acids Res. 2022;50.

66. Prieto Barja P, Pescher P, Bussotti G, Dumetz F, Imamura H, Kedra D, et al. Haplotype selection as an adaptive mechanism in the protozoan pathogen *Leishmania donovani*. Nat Ecol Evol. 2017;1.

67. Ouakad M, Vanaerschot M, Rijal S, Sundar S, Speybroeck N, Kestens L, et al. Increased metacyclogenesis of antimony-resistant *Leishmania donovani* clinical lines. Parasitology. 2011;138.

68. Van Bockstal L, Bulté D, Hendrickx S, Sadlova J, Volf P, Maes L, et al. Impact of clinically acquired miltefosine resistance by *Leishmania infantum* on mouse and sand fly infection. Int J Parasitol Drugs Drug Resist. 2020;13.

69. Hendrickx S, Van Bockstal L, Aslan H, Sadlova J, Maes L, Volf P, et al. Transmission potential of paromomycin-resistant *Leishmania infantum* and *Leishmania donovani*. Journal of Antimicrobial Chemotherapy. 2020;75.

70. LeishVet. Canine and Feline Leishmaniosis. A brief for the practicing veterinarian. 2018.

71. Baneth G, Shaw SE. Chemotherapy of canine leishmaniosis. Vet Parasitol. 2002;106:315–24.

72. Hernández L, Bolás-Fernández F, Montoya A, Checa R, Dado D, Gálvez R, et al. Unresponsiveness of Experimental Canine Leishmaniosis to a New Amphotericin B Formulation. Advances in Pharmaceutics. 2015;2015.

73. Mano C, Kongkaew A, Tippawangkosol P, Somboon P, Roytrakul S, Pescher P, et al. Amphotericin B resistance correlates with increased fitness in vitro and in vivo in *Leishmania (Mundinia) martiniquensis*. Front Microbiol. 2023;14.

74. Bryceson A. A policy for leishmaniasis with respect to the prevention and control of drug resistance. Tropical Medicine and International Health. 2001;6.

75. van Griensven J, Dorlo TP, Diro E, Costa C, Burza S. The status of combination therapy for visceral leishmaniasis: an updated review. The Lancet Infectious Diseases. 2024;24.

76. García-Hernández R, Manzano JI, Castanys S, Gamarro F. *Leishmania donovani* Develops Resistance to Drug Combinations. PLoS Negl Trop Dis. 2012;6.

77. Haldar AK, Sen P, Roy S. Use of Antimony in the Treatment of Leishmaniasis: Current Status and Future Directions. Mol Biol Int. 2011;2011.

78. Beverley SM, Coderre JA, Santi D V., Schimke RT. Unstable DNA amplifications in methotrexate resistant *Leishmania* consist of extrachromosomal circles which relocalize during stabilization. Cell. 1984;38:431–9.

79. Ubeda JM, Légaré D, Raymond F, Ouameur AA, Boisvert S, Rigault P, et al. Modulation of gene expression in drug resistant *Leishmania* is associated with gene amplification, gene deletion and chromosome aneuploidy. Genome Biol. 2008;9.

80. Downing T, Imamura H, Decuypere S, Clark TG, Coombs GH, Cotton JA, et al. Whole genome sequencing of multiple *Leishmania donovani* clinical isolates provides insights into population structure and mechanisms of drug resistance. Genome Res. 2011;21.

81. Coelho AC, Leprohon P, Ouellette M. Generation of Leishmania Hybrids by Whole Genomic DNA Transformation. PLoS Negl Trop Dis. 2012;6.

