## Additional file 3 Figure S1 for "Insights on genomic profiles of drug resistance and virulence in a cohort of *Leishmania infantum* isolates from the Mediterranean area"

**Additional File 2: Figure S1. Correlation between chromosome copy number and biomarker copy number (sub-chromosomal level).** The scatterplots show the Pearson correlation of chromosome copy number and biomarker copy number. Detection of three distinct correlation patterns: strong (extended H-locus and *LdMT),* moderate (*aqp1*, MSL loci) and low (*LdRos3*, LdSMT locus, METK locus, and PMM locus).

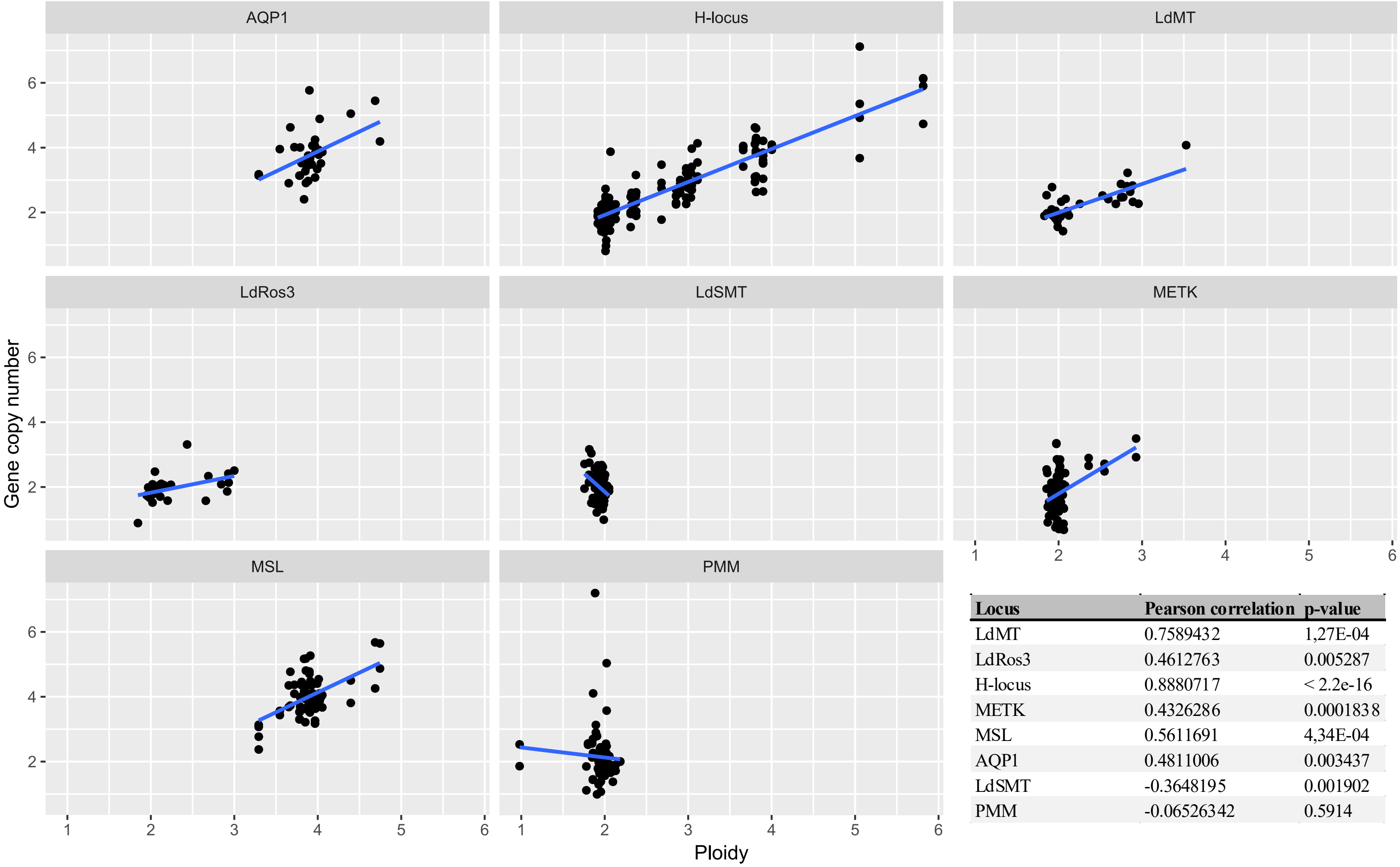


| Locus | Pearson correlation | p-value |
|---|---|---|
| LdMT | 0.7589432 | 1,27E-04 |
| LdRos3 | 0.4612763 | 0.005287 |
| H-locus | 0.8880717 | < 2.2e-16 |
| METK | 0.4326286 | 0.0001838 |
| MSL | 0.5611691 | 4,34E-04 |
| AQP1 | 0.4811006 | 0.003437 |
| LdSMT | -0.3648195 | 0.001902 |
| PMM | -0.06526342 | 0.5914 |
